# Precise MRI-Histology Coregistration of Paraffin-Embedded Tissue with Blockface Imaging

**DOI:** 10.1101/2025.06.02.657335

**Authors:** Yixin Wang, William Ho, Istvan N. Huszar, Phillip DiGiacomo, Hossein Moein Taghavi, Lee Tao, Matthew Choi, Nhu Nguyen, Samantha Leventis, David B. Camarillo, Philipp Schlömer, Markus Axer, Wei Shao, Mirabela Rusu, Inma Cobos, Jeff Nirschl, Marios Georgiadis, Michael Zeineh

## Abstract

Magnetic resonance imaging (MRI) provides 3D spatial information on tissue, yet it lacks at the molecular level. In contrast, histology provides cellular and molecular information, but it lacks the 3D spatial context and direct *in vivo* translation. Coregistering the two is key for the 3D-embedding of histological details, validating pathological MRI findings, and finding quantitative imaging biomarkers of neurodegenerative diseases. However, coregistration is challenging due to non-linear distortions of the tissue from histological processing and sectioning leading to microscopic and macroscopic nonlinear 3D deformations between specimen MRI and stained histology sections. To address this, we developed a novel pipeline, named Brewster’s Blockface Quantification (BBQ), integrating robust optical approaches with innovative 2D and 3D registration algorithms to achieve precise volumetric alignment of specimen MRI data with histological images. On a variety of brain tissue specimens from distinct anatomical regions and across multiple species, our methodology generated blockface volumes with minimal distortion and artifacts. Using these blockface volumes as an intermediary, we achieve a precise alignment between MRI and histology slides, yielding registration results with an overlapping Dice score of ∼90% for whole tissue alignment between MRI and blockface volumes, and >95% for 2D MRI-histology registration. This correlative MRI-histology pipeline with robust 2D and 3D coregistration methods promises to enhance our understanding of neurodegenerative diseases and aid the development of MRI-based disease biomarkers.

## 1. Introduction

Magnetic Resonance Imaging (MRI) provides maps of brain structure and microstructure that can noninvasively assess brain pathology in a variety of disease entities (Barthel et al., 2015; Bobinski et al., 2000; Zeineh et al., 2015; Zhang et al., 2017). However, there are limitations to MRI’s utility. For example, MRI’s current clinical role in Alzheimer’s Disease (AD) is primarily limited to excluding alternative diagnoses or evaluating for hemorrhage related to novel therapies (Hnilicova et al., 2023). Similarly, disease-specific MRI biomarkers are lacking in many neurodegenerative diseases.

One approach to developing new biomarkers is to identify direct imaging correlates of neuropathology by coupling *ex vivo* MRI with histology (De Barros et al., 2019), facilitating translation to *in vivo* human imaging. Previous studies have shown that increased iron accumulation corresponds to signal changes observed via *in vivo* MRI (Cosottini et al., 2016), which could be further evaluated using *ex vivo* ultra-high-resolution MRI for more precise pathological correlation (Kwan et al., 2012). However, achieving the required precise registration between MRI and histology is challenging due to complex 3D nonlinear deformations that occur between the two modalities. To produce histology slides, specimens are dehydrated by serial immersion in alcohol and xylene, followed by paraffin wax embedding, which introduces 3D nonlinear deformations. Subsequent microtome sectioning, slide mounting, and staining introduce additional 2D nonlinear deformations. These factors complicate the precise alignment of conventional histological images with MR data, especially since histological staining is often limited to selected slices sectioned at unknown 3D obliquities.

Current coregistration approaches consider the entire set of histological slices, reconstructing an initial histological volume to align with MRI, aiming to overcome issues of slice correspondence. Advanced reconstruction methods were developed to generate 3D histology volumes from 2D image stacks (Amunts et al., 2005; Casero et al., 2017; Johnson et al., 2010; Nir et al., 2014; Schormann et al., 1995; Yang et al., 2012). The correction of spatial distortions of the histological data heavily relies on accurate and consistent registration algorithms (Chakravarty et al., 2006; Rusu et al., 2015). For instance, Adler *et al*. designed a graph-theoretic slice stacking algorithm to correct distorted slices, followed by iterative affine and diffeomorphic co-registration with postmortem MRI scans (Adler et al., 2014). This reconstruction was further enhanced by using an interactive tool to visualize multiple histology slices with MRI (Yushkevich et al., 2016). Such approaches necessitate labor- and resource-intensive dense and uniform histology sampling and staining, and their precision is compromised by the challenges associated with sectioning angles, nonlinear slice distortions, and staining quality.

Alternative methods address this 2D to 3D transformation by optimizing the slice-to-volume registration (Gibson et al., 2012; Goubran et al., 2013; Howard et al., 2023; Huszar et al., 2023; Kim et al., 2000; Nir et al., 2014; Osechinskiy & Kruggel, 2011; Schilling et al., 2019a; Smart et al., 2023). Goubran *et al*. proposed a combined 3D and 2D registration algorithm that alternates between slice-based and volume-based registration with *ex-vivo* MRI (Goubran et al., 2013). However, there is a risk of misalignment between histology slices and 3D MRI due to 3D deformation, and the resolution difference can lead to failures during the resampling of the image volume (Pichat et al., 2018). An optical macro image was utilized to optimize the parameters of landmark-based registration of pathology and 3D-MR (Ohnishi et al., 2016). In a recent state-of-the-art method Tensor Image Registration Library (TIRL), Huszar *et al*. achieved a precise deformable registration of the sparsely sampled single-section histology images to MRI volumes of the human brain using sample photographs as intermediate space (Huszar et al., 2023). These methods are still prone to errors arising from the identification of the MRI plane corresponding to a histological slice, which is critical to achieving accurate sub-millimeter alignment between histology and MR volumes.

An alternative approach involves using blockface imaging - photographs taken during the sectioning process - as a bridge between standalone histology images and volumetric MRI data (Chakravarty et al., 2006; Dauguet et al., 2007; Groen et al., 2010; Iglesias et al., 2018; Magnain et al., 2013; Mancini et al., 2020; Schilling et al., 2019b; Uberti et al., 2009; Yelnik et al., 2007) or as a reference to reconstruct 3D histological data (Dubois et al., 2007) and then register to MRI volume in mice studies (Lebenberg et al., 2010). However, conventional blockface imaging lacks true volumetric capabilities since deeper portions of the specimen remain visible and constitute part of the reconstructed volume, complicating 3D alignment with MRI. Moreover, these methods are constrained by both 2D-to-2D and 3D-to-3D registration algorithms, as aligning volumes and slices from different modalities (MRI, histology, and blockface) with variations in thickness, angles, and resolutions presents significant challenges. Finally, face-on blockface imaging constrains the workspace for acquiring sections, limiting throughput, though some have moved the angle of the imaging apparatus to the side (Shojaii et al., 2014).

In this study, we provide a Brewster’s Blockface Quantification (BBQ) pipeline that addresses many of the above challenges. Our novel approach utilizes Brewster’s angle optics to generate high-quality and high-resolution blockface volumes that do not have depth contamination, all while providing adequate operator space. We evaluate algorithms that can compensate for nonlinear distortions occurring between MRI, blockface acquisition, and histology slide staining. We demonstrate a high-fidelity alignment between histology features and their MR counterparts, which should empower translation to *in vivo* imaging biomarkers.

## 2. Methods

### 2.1. Overview

The proposed pipeline is outlined in **Fig. 1**. After a high-resolution MRI to capture the detailed tissue structure of formalin-fixed tissue, we paraffin-embed and then serially section the entire tissue block while leveraging Brewster’s angle optics to provide accurate surface photos for each section. These photos are distortion-corrected and reconstructed into a 3D volume. We then align the MRI to the blockface volume using Tensor Image Registration Library (TIRL) (Huszar et al., 2023), providing an MRI slice corresponding to each blockface slice. Because each blockface slice/photo has a corresponding histology section, the latter can be 2D-registered to the coregistered MRI slice.

**Figure 1:**
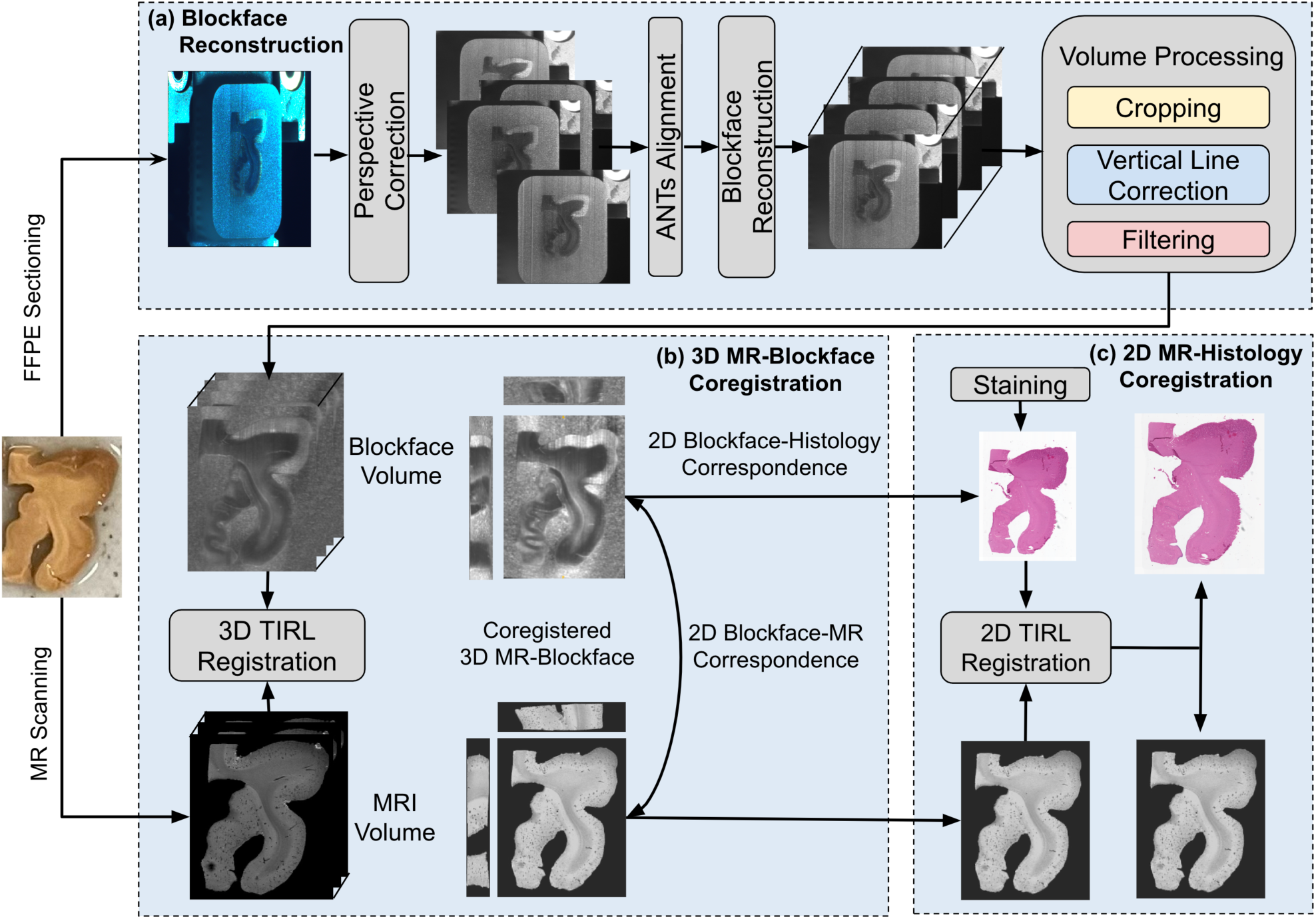
Schematic overview of image acquisition and registration pipeline. **(a)** Reconstruction of block-face volume via distortion correction, 2D alignment, and image filtering. **(b)** Determination of histology-MR slice correspondence via 3D registration of the MRI to blockface volumes. **(c)** Alignment of the derived 2D MRI slice with corresponding 2D histology stain images. Note: this 2D registration can be bidirectional.

### 2.2. Specimens

To examine generalizability across brain locations and species, we procured de-identified human brain specimens from distinct anatomical sites, namely human hippocampus and human neocortex (Stanford IRB protocol nr 33727), as well as an interior coronal slab from pig brains at the level of frontal lobe and basal ganglia (Stanford APLAC protocol nr 33684) (**Table 1**). The human hippocampal specimen, excised from a formalin-fixed human brain, was dissected, producing the hippocampal head and tail regions studied here (**Fig. 2A**). The human cortex block was dissected into multiple fragments of varying thickness, from which 2-mm and 4-mm slices were selected.

**Figure 2:**
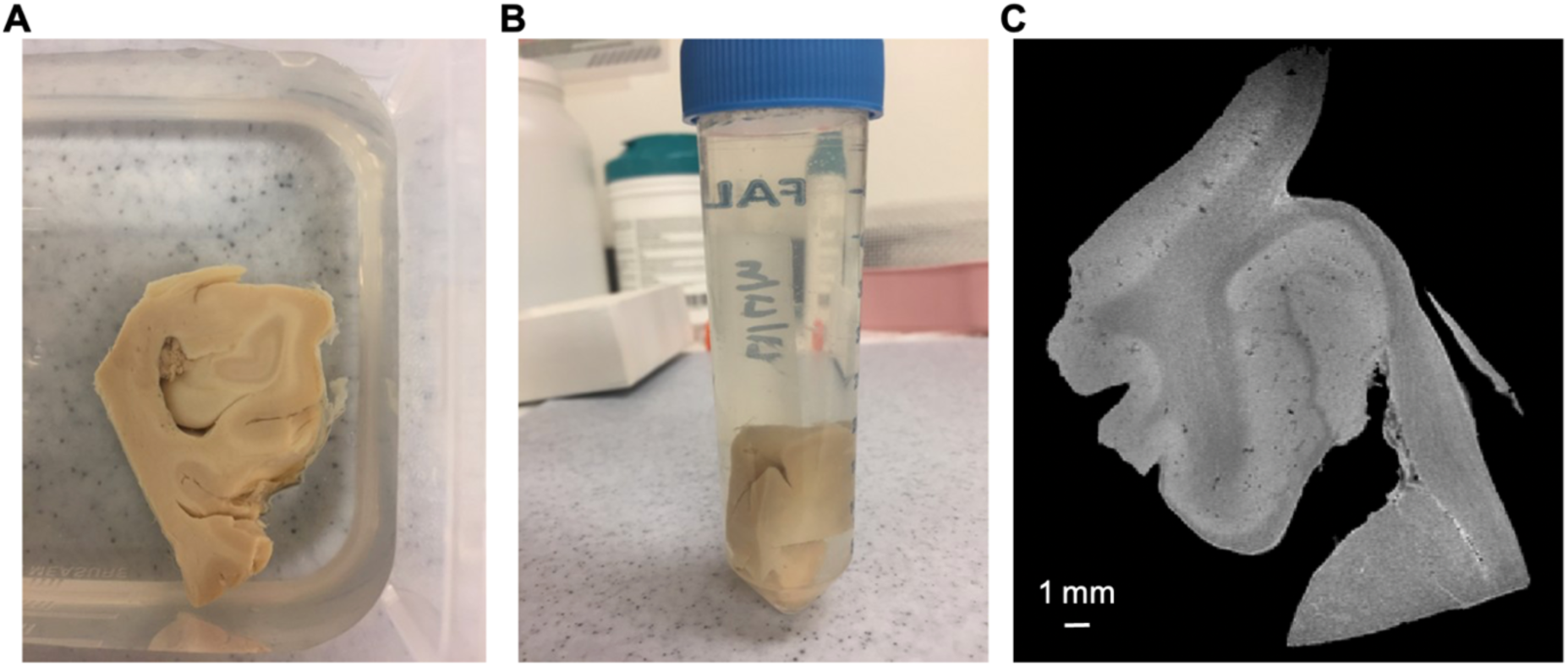
Ex-vivo specimen MRI set-up. Dissected specimens **(A)** were immersed in fluorinated oil (CHRISTO-LUBE MCG 1065) and enclosed in plastic tubes **(B)**, and then scanned using a 7T Bruker preclinical MRI **(C)**, yielding high-resolution multi-echo gradient echo (MGE) images.

**Table 1:** Overview of the five slabs used in our experiments.

| Species | Anatomic Region | Isotropic MR Resolution (mm) | Block Thickness after Formalin-fixation (mm) | Microtome Section Spacing (μm) | Blockface Thickness after Paraffin Embedding (mm) | Number of Blockface Sections |
| --- | --- | --- | --- | --- | --- | --- |
| Human | Hippocampal Head | 0.075 | 4.5 | 10 | 3.7 | 388 |
| Human | Hippocampal Tail | 0.075 | 4.2 | 10 | 3.3 | 376 |
| Human | 4-mm Cortex | 0.1 | 5.7 | 7 | 4.5 | 650 |
| Human | 2-mm Cortex | 0.1 | 2.8 | 5 | 2.4 | 545 |
| Pig | Pig Coronal Slab | 0.1 | 5.2 | 10 | 3.5 | 365 |

*Notes: (1) The thickness of formalin-fixed tissue pre-embedding is slightly smaller than the blockface thickness post-paraffin embedding due to non-linear deformation (e.g. dehydration) during tissue embedding. (2) The number of sections with the nominal slicing gap accounts to a thickness that is greater than the measured thickness of the block. This is due to the slight, unavoidable variations in the sampling angle ( $<20^\circ$ ).*

### 2.3. Image Acquisition

#### 2.3.1. MR Scanning

Formalin-fixed tissue slabs (**Fig. 2A**) were individually immersed in an inert fluorinated oil (CHRISTO-LUBE MCG 1065, ECL, Inc.) in 50ml polypropylene centrifuge tubes (**Fig. 2B**) and vacuum-degassed. We obtained high-resolution multi-echo gradient echo (MGE) *ex vivo* MRI (0.075-0.1mm isotropic) of each specimen using a 7.0T Bruker scanner using a quadrature 30 mm inner-diameter Millipede transmit/receive volume RF coil (ExtendMR LLC, Milpitas, CA). We acquired three repetitions of an MGE sequence (∼45 mins per repetition) that consisted of 10 echoes with echo times (TEs) ranging from 4-40ms (4ms separation), a repetition time (TR) of 100ms, and a bandwidth of 144kHz (**Fig. 2C**).

#### 2.3.2. Sectioning and Simultaneous Blockface Imaging

The formalin-fixed specimens underwent paraffin embedding. Special care was taken to lay the tissue within the embedding well as flat as possible, to avoid oblique sectioning of the tissue later. Paraffin blocks were mounted in a HistoCore NANOCUT R microtome (Leica, Inc.) for sectioning (**Fig. 3**). The human specimens were embedded in “clear” paraffin (Fisherbrand™ Histoplast Paraffin Wax), while the porcine specimen was embedded in “white paraffin”, a mixture of paraffin and white crayon (Ishii et al., 2021), to test different processing.

**Figure 3:**
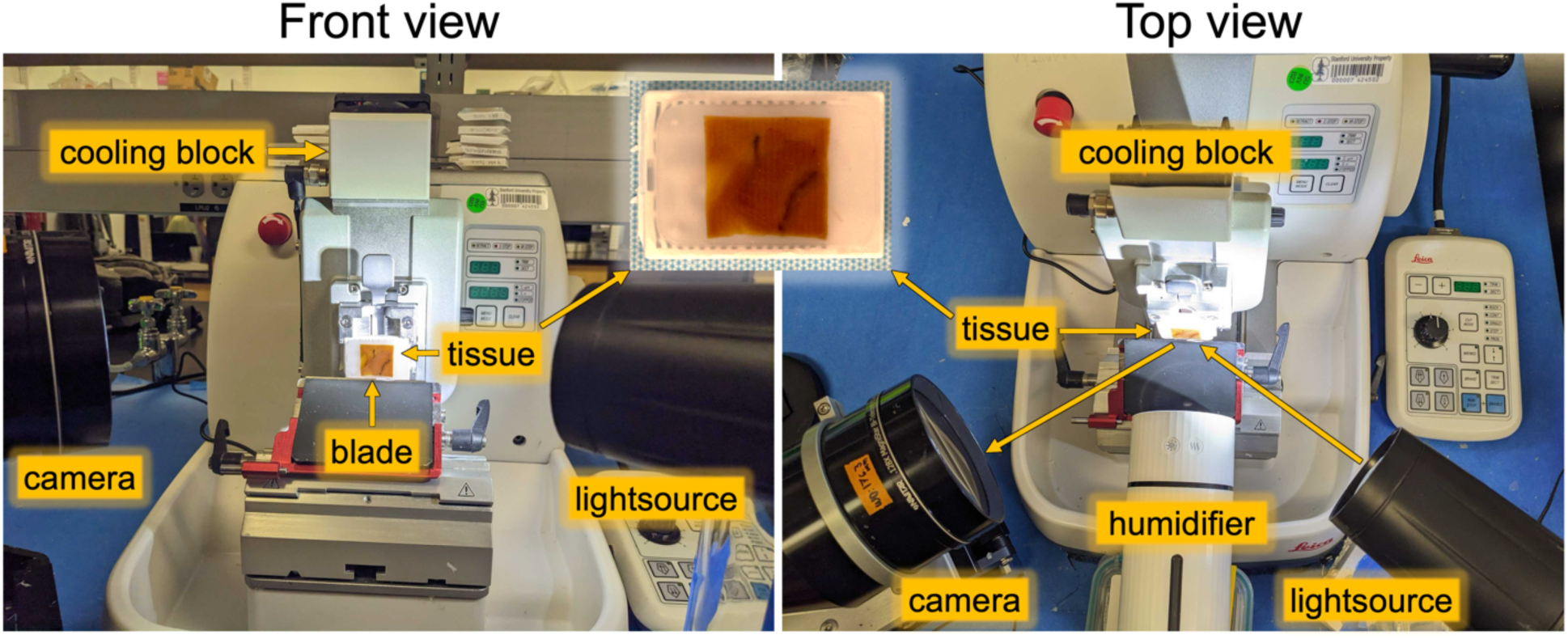
Blockface imaging setup, showing both front and top views to illustrate Brewster’s an-gle (57° to the surface normal) between lightsource, tissue, and camera. A humidifier was applied before each section was cut.

For each block, 365-650 sections were serially cut at 5-10 μm thicknesses (**Table 1**). Before each section was cut, we captured an optical image of the tissue block’s surface in the home-locked position using a 5MP color camera equipped with a bi-telecentric lens (System details: PL-D775 5.0MP rolling shutter CMOS USB3.0 color camera; Pixellink color sensor: MT9P006; mono sensor: MT9P031, Navitar, Inc., Bi-telecentric 0.0128X F/7 C-MOUNT lens, Magnification: 0.128X, Working Distance: 176.3 mm, Telecentricity: 0.05°, Field Depth: 31 mm, Average Transmittance from 460-630 nm: 97%, Navitar, Inc.) (Shojaii et al., 2014). The native in-plane resolution of each image is approximately 15 μm per pixel.

To enhance section quality, we utilized a cooling block that maintains the temperature at approximately 10°F (−12.22°C) (Leica RM CoolClamp, Leica, Inc.) on the microtome to prevent the paraffin block from heating and softening, achieving more uniform sectioning. Additionally, a rechargeable portable humidifier (Palanchy, Inc.) was pointed at the tissue for approximately 5 seconds before each cut to prevent tissue from rolling up and to maintain section quality.

The goal of blockface imaging is to capture the undistorted state of the surface of the block prior to cutting each section. Challenges include visualizing just the surface of the block without imaging deeper tissue, providing operator workspace in front of the tissue, and avoiding image distortion. To surmount these challenges, we position both the camera and our LED lightsource at Brewster’s angle (57° from the normal for the paraffin-air boundary) (**Fig. 3, Top view**). Brewster’s angle is where only the S-polarized light (perpendicular to the plane of incidence) is reflected, while P-polarized light (parallel to the plane of incidence) is transmitted through the surface, thus eliminating depth information (Brewster, 1815; Hui, 2020). A linear polarizer sheet was positioned in front of the camera to allow only the reflected light, containing purely surface information, to reach the camera. Adequate workspace is available when using this angle. To minimize perspective distortion, we use a telecentric lens (**Fig. 3**) and acquire calibration images of a printed grid of known dimensions before each cutting session (**Fig. 5**). (See Supplementary Protocols for detailed setup descriptions).

We observed that illuminating the tissue precisely at Brewster’s angle not only resulted in low tissue contrast (**Fig 4, top**), but also became overly sensitive to surface irregularities. In particular, we observed vertical line artifacts that were most likely associated with the motion of the blade through the heterogeneous material composition of the block. Therefore, we adjusted the position of the lightsource to be slightly off Brewster’s angle (See Supplementary Protocols), (**Fig. 4, bottom**). This angle of illumination (hereafter referred to as “Brewster’s adjacent”) was further optimized to maximize tissue contrast and minimize the visibility of tissues beneath the surface via raising the light source by a few degrees and angling it downward. In combination with our polarization filter, this resulted in high tissue contrast and effective imaging of the tissue surface. To maximize the homogeneity of this angle, the tissue block was positioned with its long axis horizontal (**Fig. 3**). We note that moving too far off Brewster’s angle eliminated the vertical-line artifact but introduced undesirable contaminating depth information. We also noted that gray-white contrast was inverted on human tissue between Brewster’s angle vs. Brewster’s Adjacent.

**Figure 4:**
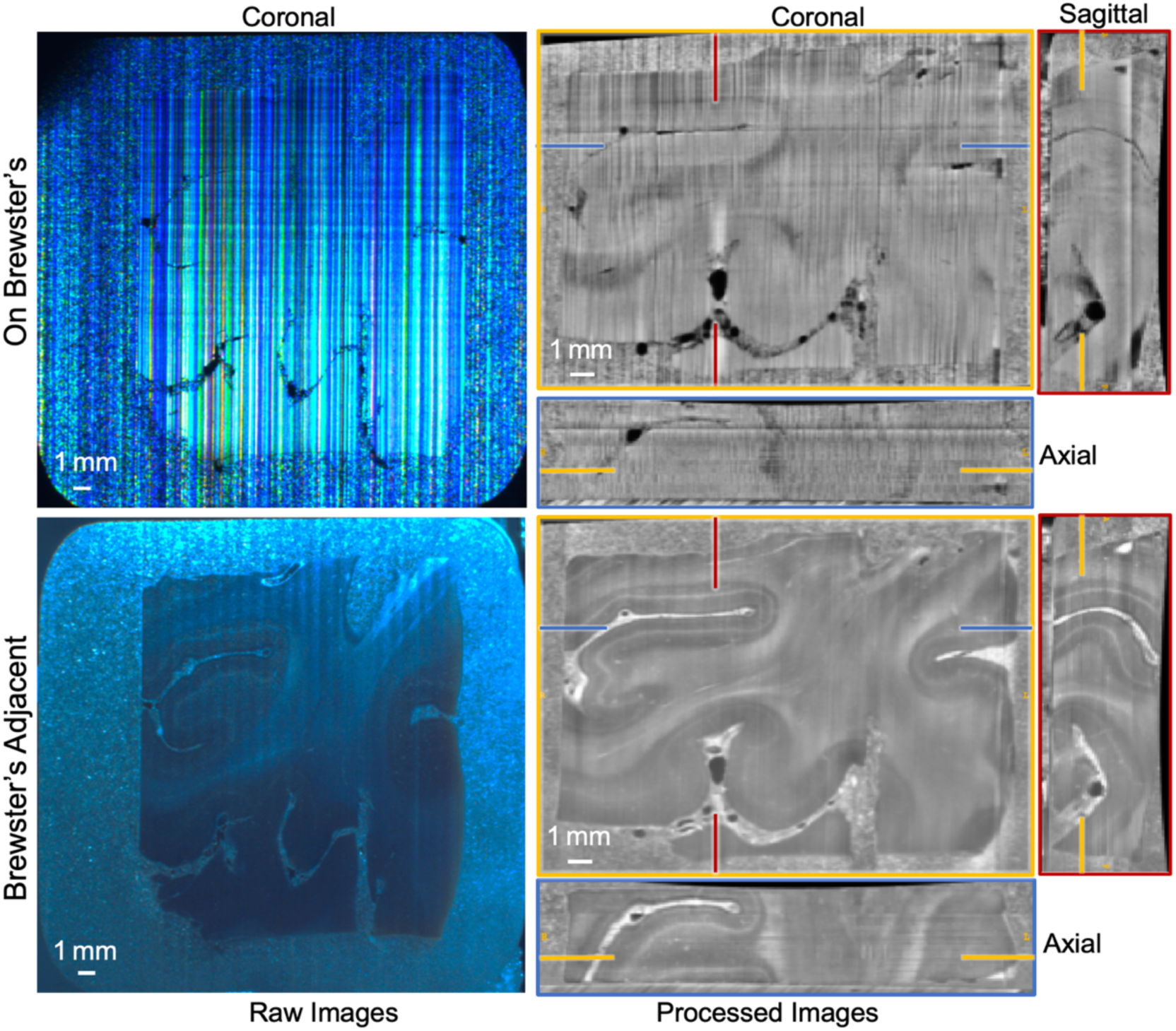
Comparison of simultaneously acquired raw and processed blockface images of a neocortical human brain specimen captured under two different lighting setups: (**top row**) directly at Brewster’s angle (On Brewster’s) and (**bottom row**) slightly off Brewster (Brewster’s Adjacent). Note the inversion of gray-white contrast between the two positions.

In addition to the imaging setup described above, we also experimented with a front-facing camera and used white paraffin to eliminate depth information. In our experiments with the pig specimen, this approach reduced structural details and edge definition and provided less room for the operator to collect the slices (**Supplement Fig. S1**).

#### 2.3.3. Histology

During serial sectioning of the blocks, we sparsely captured sections and placed them in a conventional water bath, then mounted each onto charged slides. Slides underwent staining with hematoxylin and eosin (H&E) and imaging at a Leica AT2 whole-slide scanner (magnification: 20X, pixel size: 0.5 μm) using ImageScope software (v12.4.3.5008). Labeling histology slides with the corresponding blockface image number established a precise one-to-one correspondence between histological sections and respective blockface images. For efficient processing and registration, we utilized histology images from Series 2 (∼3000 × 3000 pixels), corresponding to a resolution of approximately 8 µm per pixel. For the 2-mm block, we stained and slide-scanned 19 sections (approximately every 100 μm throughout the block to match MR resolution) and created a high-resolution histological volume.

### 2.4. Image Processing

#### 2.4.1. MRI

To reduce image noise, we concatenated all repetitions (3x), echoes (10x), and real-imaginary coils (2x), resulting in a total of 60 volumes, and denoised using MP-PCA (Veraart et al., 2016) (*dwidenoise* from MRtrix3^1^), which resulted in small incremental improvement compared with averaging all magnitude images. Afterward, we unmerged the denoised volumes, computed magnitude images, registered all repetitions by transforming the second and third repetition to the space of the first (to account for minor translations that occur during scanning), and removed ringing artifacts using MRtrix3’s *mrdegibbs* (Kellner et al., 2016). Finally, the data from the echoes were averaged to obtain a single final image with higher contrast and higher signal-to-noise ratio (**Fig. 2c**).

#### 2.4.2. Blockface Images to 3D Volume

##### Perspective correction

The telecentric lens resolved perspective distortion in the vertical direction so the paraffin block did not get smaller to the right where it is more distant (**Fig. 5b**). Yet, there was still aspect ratio distortion along the left-right axis. To rectify this, we used the same setup to capture photographs of a fixed grid image with dots of known spacing of 5 millimeters. These images were positioned at the same location as the tissue cutting plane in the microtome (**Fig. 5a**). An in-house MATLAB script semi-automatically (i.e., using user’s input in a MATLAB color-thresholding user interface) segmented the grid’s intersection points and computed an affine transformation matrix between this grid image and its known dimensions. This matrix was then sequentially applied to each 2D image to correct distortion. This calibration should be identical for different specimens assuming the constant position of the optics relative to the microtome, though we always collected these images prior to and/or at the conclusion of sectioning.

**Figure 5:**
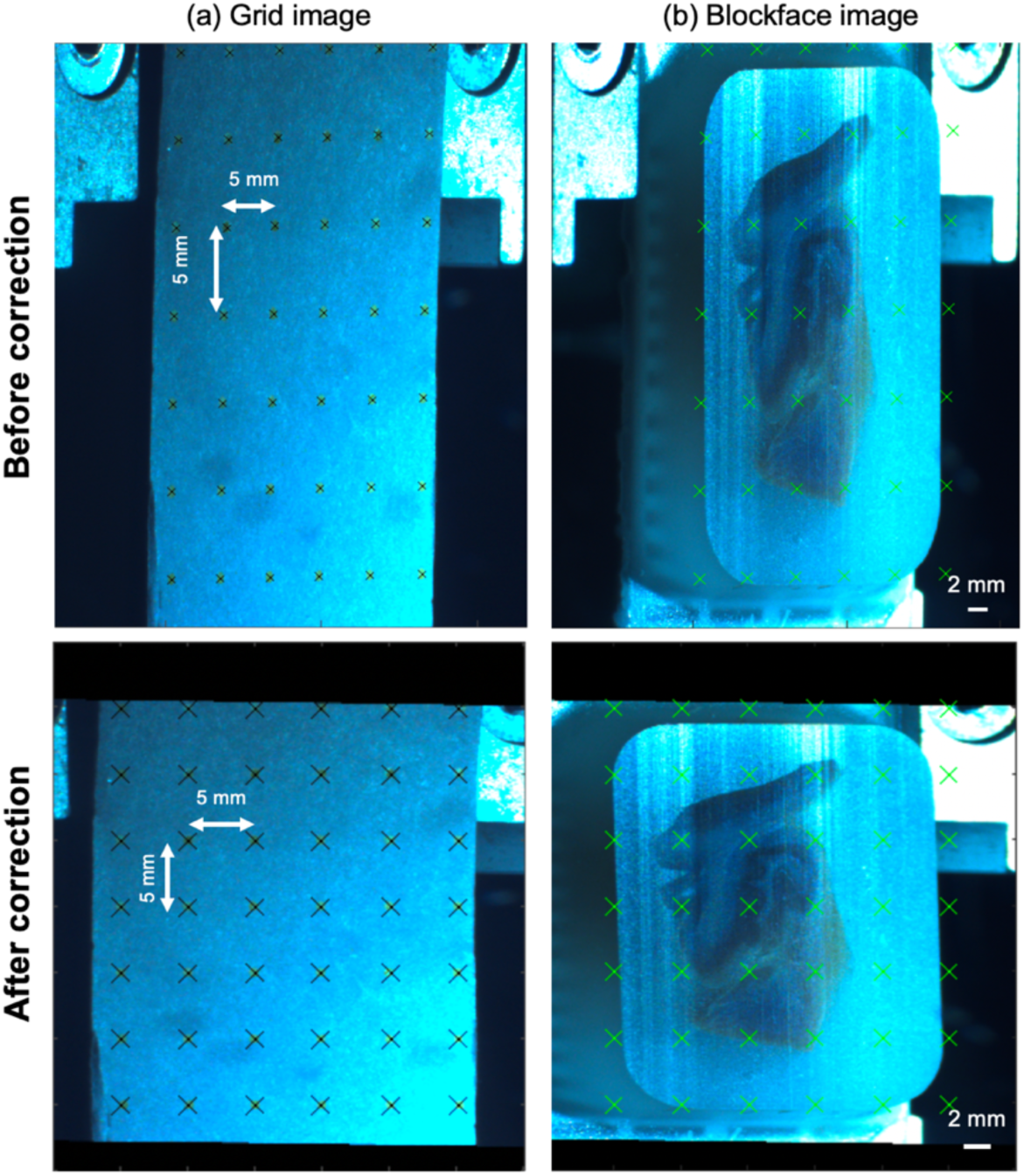
Blockface perspective correction. (**a**) A fixed grid, printed and adhered to the micro-tome mount, was imaged (first row) and transformed to the correct perspective without distortion (second row). (**b**) The same transformation matrix was then applied to the blockface images captured from the identical position.

##### Quality control, 1D translation correction, and cropping

Before constructing a blockface volume, individual images were visually inspected. Rare low-quality images (e.g., the operator’s hand was in front of the block, or the image was captured while the microtome was still moving) were manually identified and excluded. For each excluded image, a duplicate of the previous slice was added to maintain sequence continuity. The home position for the microtome has limited precision. To account for this and other minor differences in positioning from section to section, we utilized Advanced Normalization Tools (ANTs) (Avants et al., 2009) for aligning serial 2D blockface images, only correcting for 1D vertical translation.

To compose the 3D volume, each blockface image was cropped to isolate the paraffin block (**Supplement Fig. S2(a)**) and resampled at an isotropic in-plane resolution of 0.0125-0.025 mm/pixel, and image sizes of approximately 35 mm x 30 mm (approximately 2,000 pixels by 1,000 pixels).

##### Artifact removal

Even in the Brewster’s adjacent imaging configuration, vertical line artifacts are present in the images, prompting us to devise two correction techniques.

###### Technique A: Moving Average Method

A predefined sliding window iteratively traversed the image columns, calculating the average intensity of pixels in the neighboring columns on either side of the center column within the window. This process normalized each pixel in the center column, effectively smoothing abrupt intensity variations and diminishing the prominence of vertical line artifacts, as shown in **Supplement Fig. S2(c)**. Subsequently, median filtering was applied to the entire volume, serving to further reduce noise and enhance the clarity of the images (**Supplement Fig. S2(c)**).

###### Technique B: Fourier Domain Correction (Supplement Fig. S2(d)

Initially, median filtering was applied across the entire volume as a preparatory step to reduce noise. Subsequently, each 2D slice underwent Fast Fourier Transform (FFT). The vertical lines in the spatial domain appeared as horizontal lines in the frequency domain (**Supplement Fig. S2(e),** orange arrows), which we automatically masked out, not modifying the center of the frequency domain image (**Supplement Fig. S2(f),** orange arrows).

The higher-quality results from the two techniques were selected (Technique A for the four human specimens and Technique B for the pig coronal slab) and used for subsequent MRI coregistration.

### 2.5. Coregistration

#### 2.5.1. 3D MR-Blockface Coregistration

Before automated coregistration, we manually reoriented both MRI and reconstructed blockface volumes to roughly match their orientation using ITK-SNAP (Yushkevich et al., 2006). Subsequently, a deformable, invertible registration was performed utilizing the TIRL method (Huszar et al., 2023), aiming at transforming MRI volumes to the blockface space. TIRL translates pixel/voxel coordinates into physical coordinates through a meticulously organized transformation chain, bifurcated into internal components for preserving image resolution and external components for facilitating user-driven optimization processes. TIRL accomplished blockface-MR registration by optimizing serial 3D rigid, isotropic scaling, affine, and nonlinear transformations.

To bridge the substantial domain gap between the two image modalities and quantify their similarity, the Modality Independent Neighbourhood Descriptor (MIND) (Heinrich et al., 2012) is employed within TIRL. The computation of MIND involves operating on patches around any location *x* in an image (*I*). A patch consists of the immediate neighbors of the center pixel/voxel, and patch-wise distances (*D_p_*) are calculated as a sum of squared intensity differences between corresponding pairs of pixels/voxels between two patches. By using MIND, an image will be represented by a vector at each location *x*, which is formulated as:

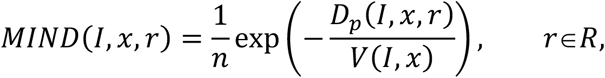

where *D_p_*(*I*, *x*, *r*) represents patch-wise distances between locations *x* and *r* in the image *I*, *V* denotes a patch-wise variance estimate at location *x*, and *R* is a spatial search region around location *x*, with *n* serving as a normalization constant ensuring a maximum value of 1. MIND representations of the images are typically calculated many times during registration for each modality, at multiple resolution scales. The search region *R* is usually defined as the immediate or an extended neighborhood around *x*. Using the immediate neighborhood definition, 3D images are represented with a six-dimensional vector at each voxel, which describes the intensity relationships of the voxel with its six immediate neighbors. As such, MIND is sensitive to intensity gradients and can describe the unique structural characteristics of an image, such as points, edges, and homogeneous areas, independently of the modality, contrast, and noise levels. This makes it particularly suitable for comparing images from MRI and blockface volumes.

The registration is driven by minimizing a cost function that is calculated as a sum of pixel-wise Euclidean distances between the MIND vectors of the MRI and blockface volumes. Additional regularization (penalizing large local deformations) is used to ensure that non-linear transformations remain smooth and invertible. This cost function was minimized in four steps, by successively optimizing the parameters of the aforementioned three linear and one non-linear transformation (the configuration file is in our online open-source code)

For comparison purposes, we performed a similar invertible 3D alignment with the commonly used ANTs Rigid, Affine, and Nonlinear transforms (Rigid: shrink factors: 16×8×4×2, gradient step: 0.01, similarity metric: mutual information; Affine: shrink factors:16×8×4×2, gradient step: 0.1, similarity metric: mutual information; and Symmetric Normalization (SyN) transformation: shrink factors: 32×16×8×4, gradient step: 0.1, smoothing standard deviation: 3 and similarity metric: cross-correlation). Affine used the Rigid output as an initial estimate, and SyN utilized the Affine output. To ensure a fair comparison, we optimized key parameters through preliminary experiments on the hippocampal specimens (similarity metrics: mutual information vs. cross-correlation; gradient steps: 0.01, 0.05, 0.1, 0.15). We manually initialized the registration (rotation and scaling) using ITK-SNAP to align the two volumes as accurately as possible before applying ANTs. Free-form deformation registration (Rueckert et al., 1999) used in previous works (Dubois et al., 2007; Lebenberg et al., 2010) was also explored as a nonlinear alternative in **Supplement Fig. S3**.

#### 2.5.2. 2D MR-Histology Coregistration

After transforming the MR volume to the blockface space, we extracted the specific 2D MRI slice that corresponds to a single blockface image, thus also corresponding to a single stained histology slide. Subsequently, we executed a 2D invertible registration between each MRI and histology slide using TIRL with MIND as a cost function which encompassed 2D Rigid, 2D Affine, and 2D Nonlinear transformations. This registration is bidirectional depending on the downstream goal: when reconstructing volumetric histology of the 2-mm human cortex in Section 2.5.3, we registered Histology to MR to maintain spatial consistency across slices and align with the 3D MRI coordinate system. This can also be done at histology resolution. However, if the goal is to map MRI data to a single or isolated high-resolution histological slice or to correlate MRI images with precise histological markers or pathology at histology resolution, the histology space would serve as the target space (the rest of the specimens used for testing this purpose).

For comparison purposes, we performed the same 2D alignment using ANTs Rigid, Affine, and Nonlinear transformation (Rigid: shrink factors: 16×8, gradient step: 0.1, similarity metric: mutual information; Affine: shrink factors:16×8, gradient step: 0.1, similarity metric: mutual information; SyN: shrink factors: 16×8, gradient step: 0.05, smoothing standard deviation: 3 and similarity metric: cross-correlation). The parameters optimization for 2D alignment followed those used in the 3D registration described in Section 2.5.1, though no manual initialization was required for the 2D registration process. Free-form deformation registration was also explored as a nonlinear alternative in **Supplement Fig. S4**.

#### 2.5.3. 3D MR-Volumetric Histology

It is possible within an MRI-histology registration framework that an apparent high-fidelity alignment seen between the final 2D images is not representative of true 3D geometry but is instead driven by an excessive nonlinear transformation, especially of the final 2D to 2D registration. To assess the volumetric registration in 3D, the 19 uniformly sampled histology images from the 2-mm human cortex were coregistered to their 19 corresponding 2D MR slices and reconstructed into a 3D histology volume in the space of the MR volume.

### 2.6. Registration Accuracy Quantification

A segmentation of MRI, blockface, and histology images for three specimens was performed manually in each modality’s native space, including 3D MRI, 3D blockface, and 2D histology, using ITK-SNAP (Yushkevich et al., 2006) in order to assess registration accuracy. Boundaries were sometimes easier to see in MRI (i.e. the MRI better defined the grey-white boundary than the blockface images), so native-space MRIs were sometimes utilized as a reference for rough structural localization for blockface to avoid structural ambiguity. All manual segmentations were completed prior to any registration and therefore were not influenced by the registration outcome. Histological images were segmented without referring any other modalities.

We segmented: (1) the human hippocampal regions’ white matter, grey matter, and dentate gyrus, (2) the human cortex white matter and grey matter, and (3) the pig white matter, grey matter, caudate, putamen, and anterior commissure.

Using the above 2D/3D registrations, we transformed the segmentations into the same target space using nearest-neighbor interpolation and calculated the Dice similarity coefficient to quantify overlap (Dice, 1945): 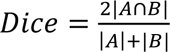, where A/B=voxels in the segmentations being compared, with a higher Dice score indicating better registration accuracy. The Dice score was calculated for each anatomical label individually and for the overall tissue by combining all labels.

For 3D MR-Volumetric Histology, in addition to quantifying segmentation overlap between MRI, blockface volumes, and histology, several vessels were manually segmented separately on both MRI and histology volumes and rendered using 3D Slicer (Fedorov et al., 2012) for 3D visualization of internal structures. We extracted skeleton structures from the segmentation masks of vessels and uniformly resampled parametric curves fit to the skeleton to spatially assess their detailed alignment in 3D.

## 3. Results

### 3.1. 3D MR-Blockface Coregistration

Visualization and segmentation of blockface and coregistered MR images (**Fig. 6**) qualitatively demonstrated the correspondence in overall tissue shape and boundaries. Overlap/discrepancies between red outlines (blockface segmentation) and blue outlines (MRI segmentation) indicated the overlap after registration (**Fig. 6 (c, d)**). Both ANTs and TIRL visually performed well in the hippocampal head and human 4-mm cortex, but TIRL achieved tissue boundaries with more overlap (orange arrows in all three planes). Consistent with this, TIRL proved qualitatively superior for the hippocampal tail and pig tissue, particularly at the dentate gyrus and anterior commissure (orange arrows in coronal planes).

**Figure 6:**
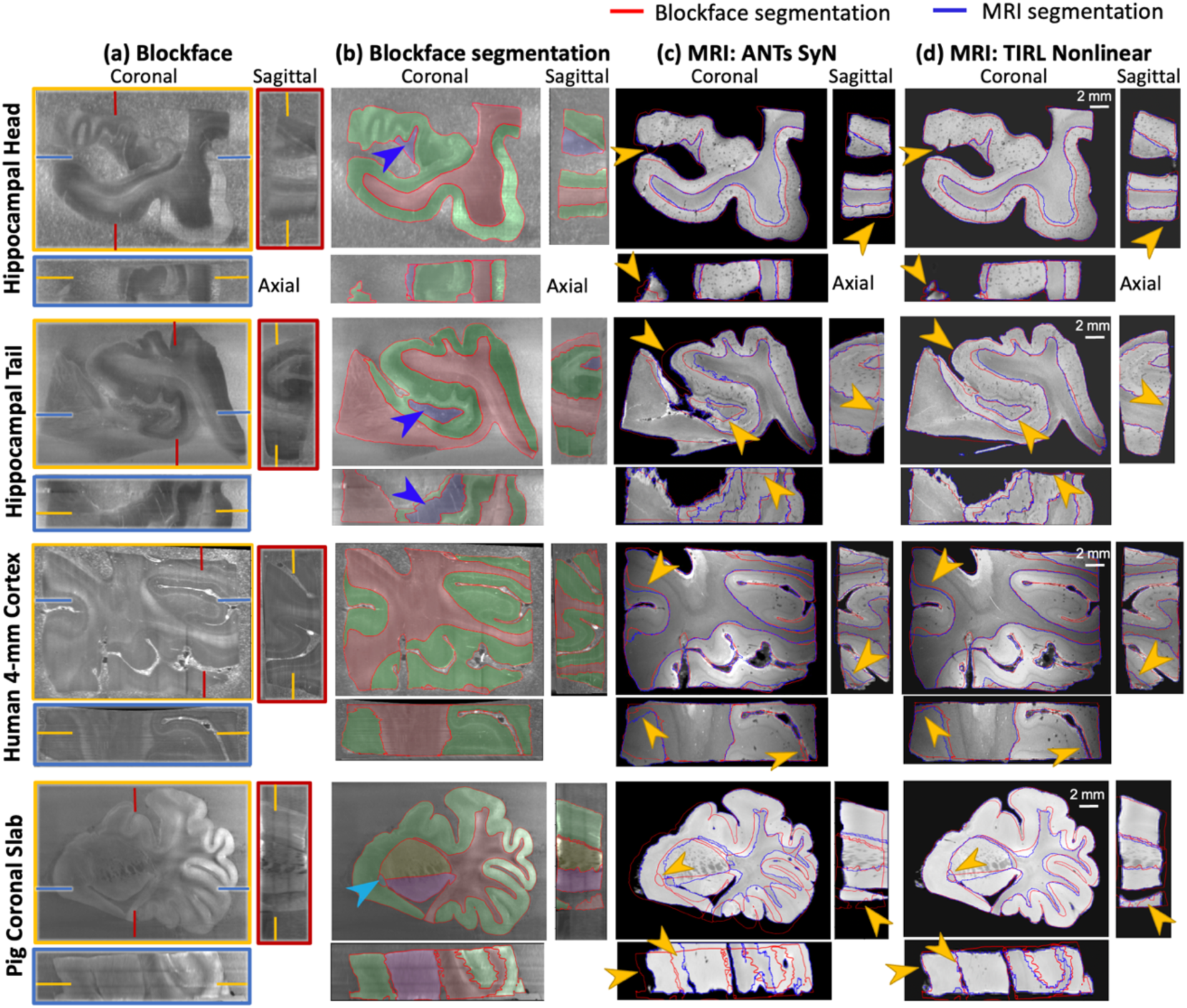
3D coregistration between MRI and blockface volumes. (**a**) Blockface images in axial, sagittal, and coronal planes. (**b**) The same sections with manually segmented for all specimens: white matter (red), grey matter (green), for hippocampal specimens: the dentate gyrus (blue, arrow), and only for pig: caudate (yellow), putamen (purple), and the anterior commissure (light blue, arrow). Red contours mark segmented region boundaries. (**c, d**) MR images after deformable (non-linear) transformation into the blockface space. Red and blue outlines trace the boundaries between subregion segmentations performed on the blockface and MRI respectively.

Quantitatively, the Dice scores (**Fig. 7**) showed the pipeline produces high overlap scores in all larger regions and most smaller regions. TIRL outperformed ANTs in all six regions and significantly outperformed ANTs Nonlinear on white matter, grey matter and whole tissue (*p=0.043*). including small but important regions such as the dentate gyri in the hippocampal head, hippocampal tail (**Fig.6(b)**, blue arrow) and pig anterior commissure (**Fig.6(b)**, light blue arrow), achieving 77.6%, 83.7% and 65.8% Dice scores, respectively. In contrast, ANTs Nonlinear (SyN) failed to align any tissue voxels in the anterior commissure. The one area ANTs performed slightly better was in the whole pig coronal slab outline, though TIRL outperformed on all smaller regions.

**Figure 7:**
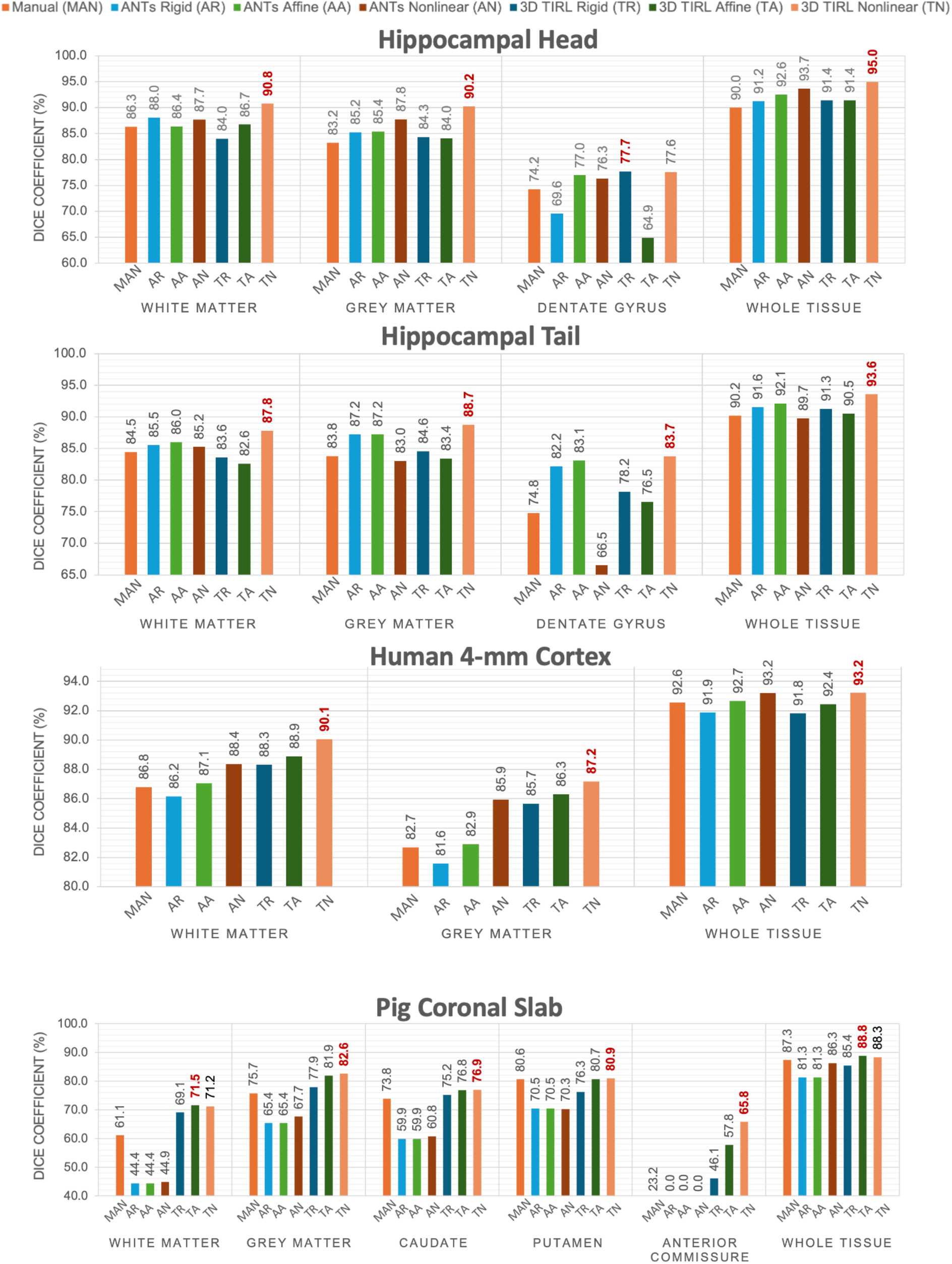
Dice similarity coefficient scores (%) of the segmentations of MRI and blockface volumes after coregistration across different specimens. “Whole Tissue” represents the score for the entire composite area. The highest score (best overlap) is indicated in **Red**.

### 3.2. 2D MR-Histology Coregistration

Figure 8 presents a qualitative assessment of the two-dimensional coregistration between histological sections and MRI images of the four specimens. Overlapping segmentations are displayed in both histology and MRI space, using transformations provided by ANTs Nonlinear (SyN) (**Fig. 8(ii)**) or TIRL Nonlinear (**Fig. 8(iii)**). For all four specimens, we obtained consistent tissue alignment despite typical tissue distortions. We found that TIRL enabled a closer correspondence of the tissue boundaries and outperformed ANTs in the dentate gyrus (orange arrows, first two rows), white-grey matter junction (orange arrows, human 4-mm cortex), and in the caudate, putamen, and anterior commissure (orange arrows, pig coronal slab).

**Figure 8:**
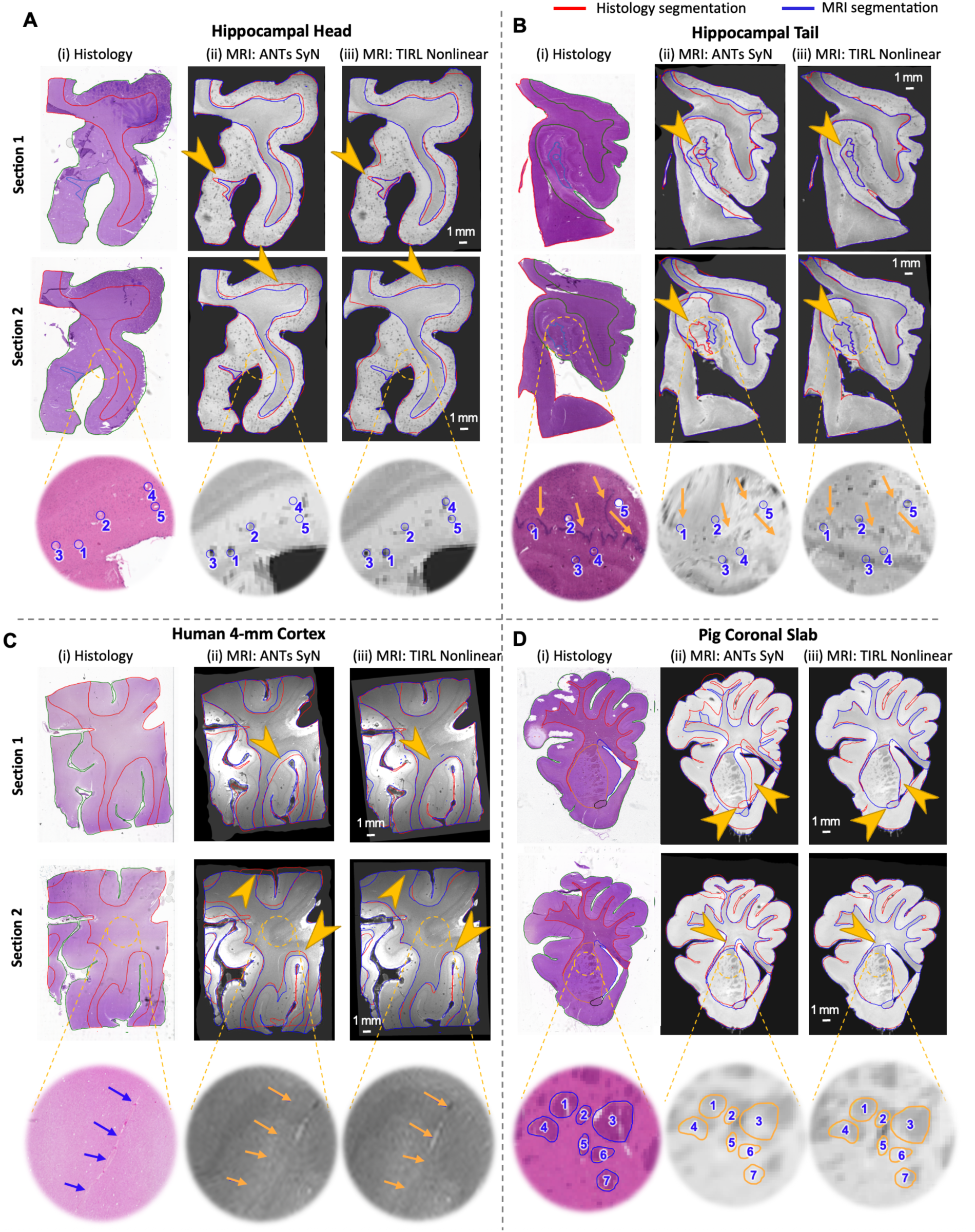
Qualitative evaluation of 2D coregistration between histology and MRI correspondences. **(i)** displays H&E-stained sections with manually labeled outlines of white matter (red), grey matter (green), dentate gyrus (light blue), caudate (yellow), putamen (purple), and the anterior commissure (black). **(ii)** and **(iii)** present MRI images transformed into the histology space. Manually annotated regions of interest within the circles and pointed by arrows on the histology slides correspond precisely to MRI images transformed using TIRL Nonlinear registration.

We achieved nearly micron-level accurate registration, evident in TIRL registrations of minute anatomical structures such as focal hypointensities representing vessels or perivascular spaces (**Fig. 8A,B**, bottom, blue circles), the dentate gyrus granule cell layer (**Fig. 8B**, bottom, orange arrows), vessels (**Fig. 8C**, bottom, orange arrows) and the internal capsule interwoven between basal ganglia nuclei (**Fig. 8D**, bottom, circles). This was quantitatively demonstrated in **Fig. 9**, where both TIRL Nonlinear and ANTs Nonlinear (SyN) achieved high Dice scores (>90%) for white and grey matter, but TIRL Nonlinear significantly outperformed ANTs SyN in white matter (*p < 0.001*), grey matter (*p = 0.002*) and whole tissue (*p = 0.041*) across all sections. TIRL Nonlinear also achieved remarkably higher mean Dice scores for finer structures such as the Dentate Gyrus and Anterior Commissure (AC) across all sections.

**Figure 9:**
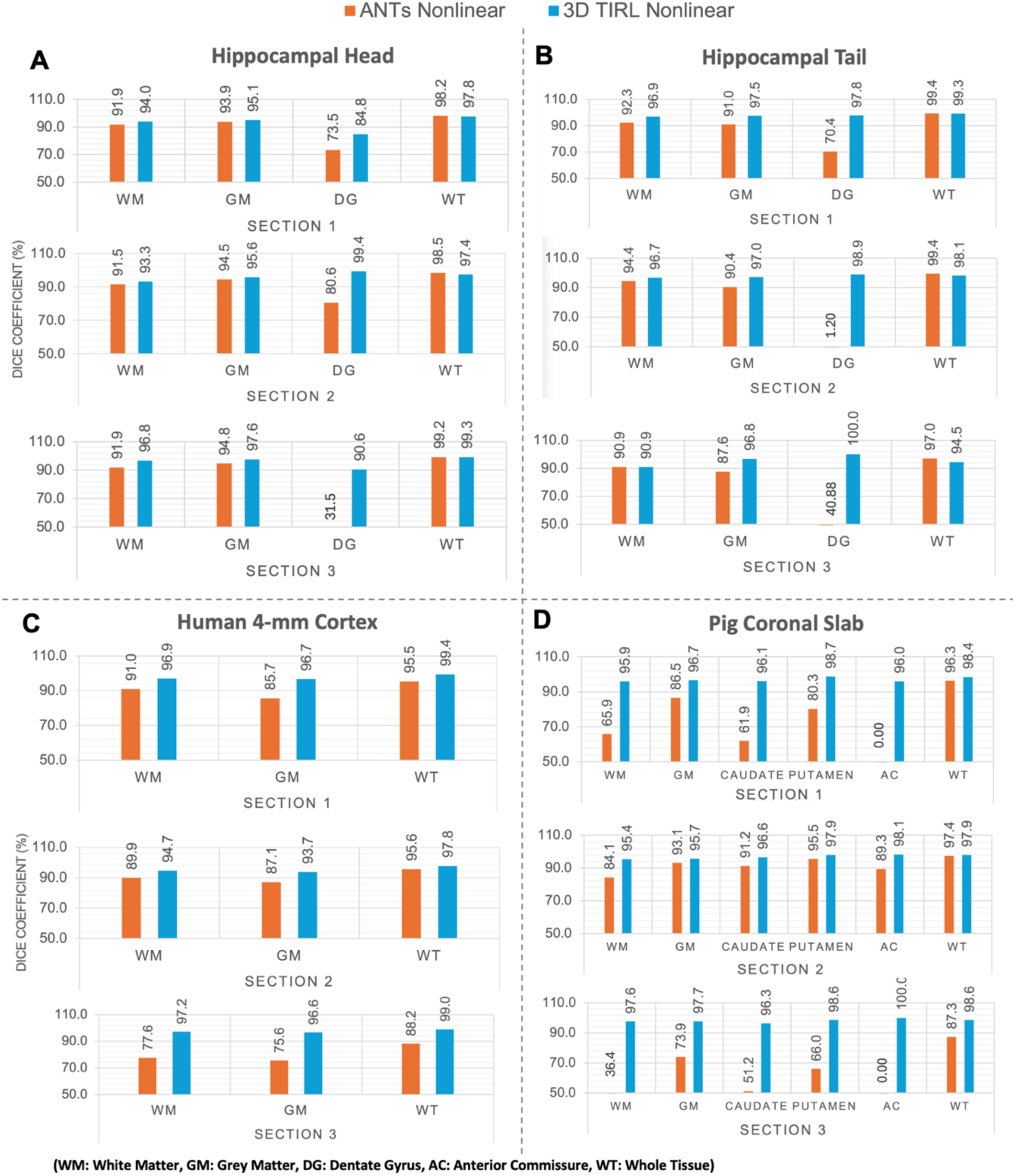
Dice similarity coefficient (%) analysis for 2D registration of MRI and histology across multiple hippocampal head, tail, 4-mm cortex and pig coronal slab slides with H&E staining. TIRL shows a superior performance in the dentate gyrus (DG) and Anterior Commissure (AC).

### 3.3. 3D Volumetric Histology

In order to ensure that the precise registrations demonstrated thus far reflect true 3D alignment of tissue structures rather than simply overfitting a 2D nonlinear registration, we used the pipeline for registration of the human 2-mm cortex MRI with uniformly sampled volumetric histology staining to produce a histology volume coregistered with MRI (**Figs. 10-11**). Blockface-MRI registration was again best achieved with TIRL Non-linear (**Fig. 10A**), which outperformed ANTs Nonlinear, as further confirmed quantitatively in **Fig. 10B**. Registration between the extracted 2D MRI slices and corresponding stained sections of all 19 uniformly sampled slices across the entire tissue (**Fig. 11**), demonstrating a Dice score of over 90% for most slices with 2D TIRL, with lower scores at slices near the edges with incomplete tissue. The resultant histological volume showed an expected correspondence when viewed from multiple orientations and in 3D, closely matching MRI (**Fig.12A/B**). To quantitatively demonstrate the 3D nature of the alignment, we separately manually segmented vessels/perivascular spaces that spanned all 19 serial slices from each modality. Comparing the segmentations, they show a visually similar shape of the structures for both modalities (**Fig.12C**). Quantitative comparison of two of these vessels (yellow and blue) showed a high degree of overlap (**Fig.12D**), with minimal non-overlapping regions (red from MRI and green from histology) and a mean Euclidean distance of 0.08mm and 0.11mm (**Fig.12E**), at or even below MRI resolution.

**Figure 10:**
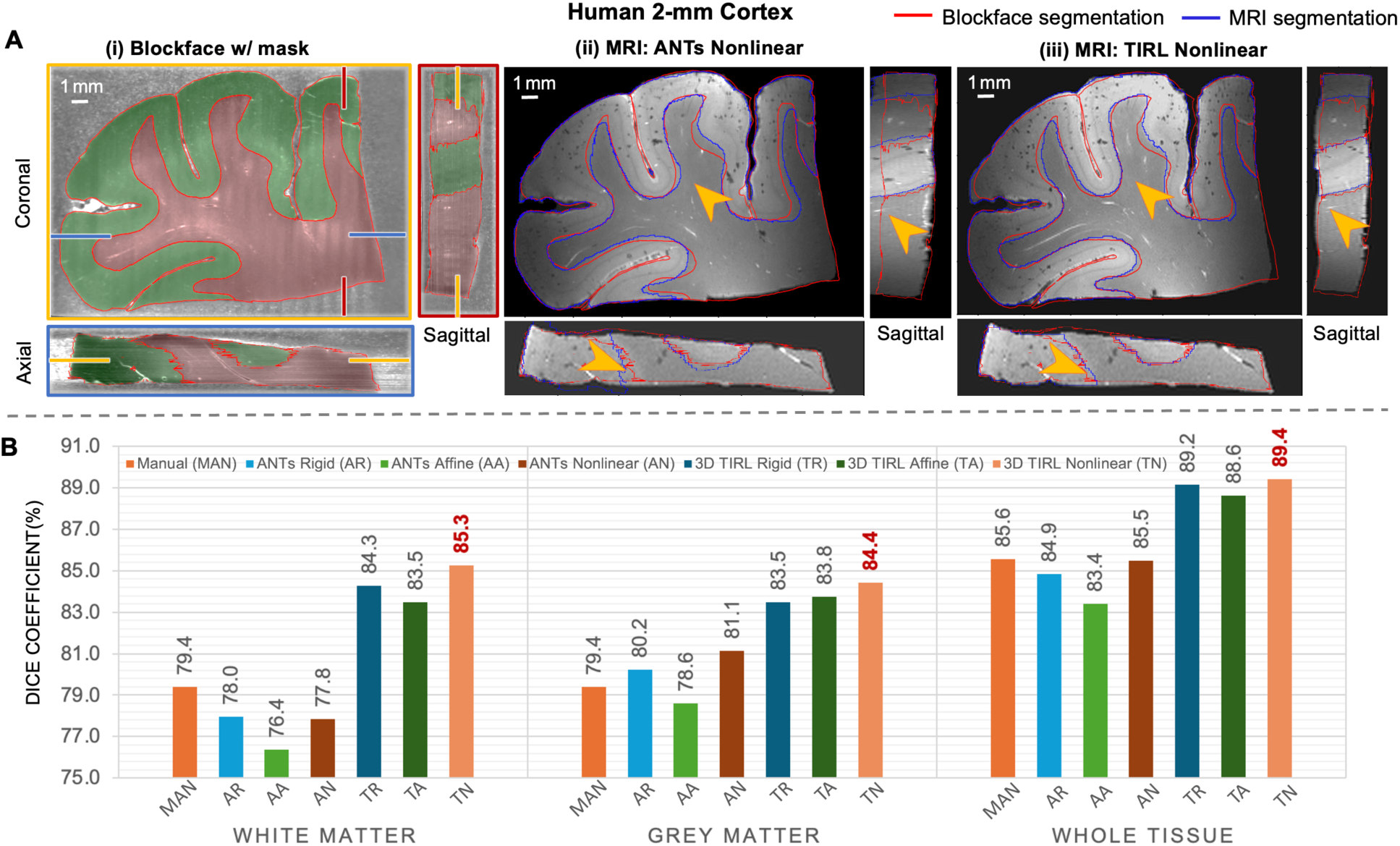
3D coregistration between MRI and blockface volumes of Human 2-mm Cortex. **A(i).** Original blockface images in 3D planes with manually segmented white matter (red), grey matter (green), and red contours marking segmented region boundaries. **A(ii, iii).** MR images after undergoing deformable (non-linear) transformation into the blockface space using ANTs Nonlinear (SyN) and TIRL Nonlinear. **B** presents Dice coefficient across subregions using different registration algorithms.

**Figure 11:**
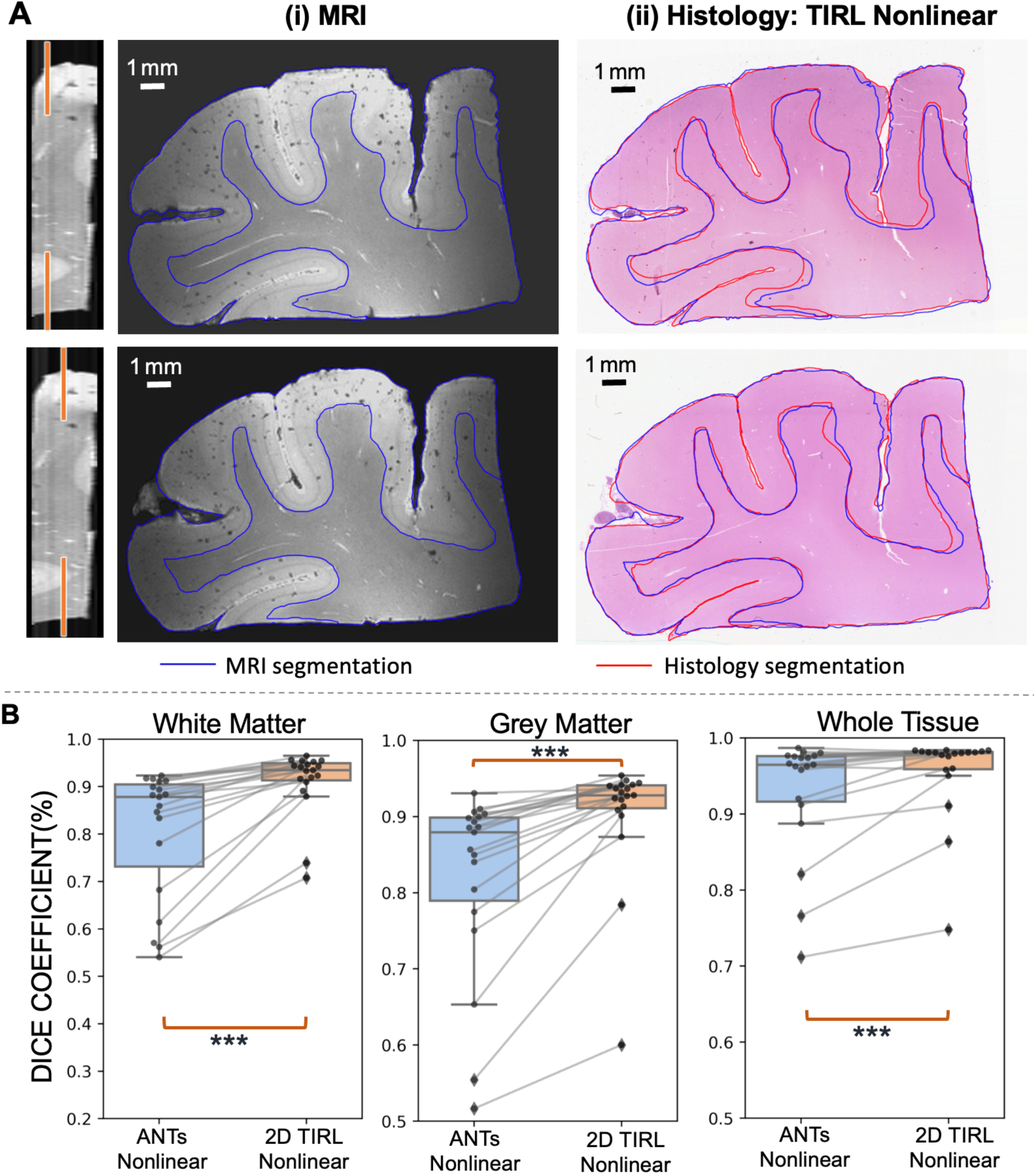
2D coregistration between MR and histology correspondences of Human 2-mm Cortex. **A(i)** displays manual segmentation of white matter and grey matter on MRI slices, and **A(ii)** presents the corresponding histology sections transformed into MRI space using TIRL Nonlinear. **B** shows the quantitative Dice coefficient for each coregistered histology slide to its corresponding MR slice using TIRL Nonlinear and ANTs Nonlinear (***: p < 0.001). Each dot represents a single slice.

**Figure 12:**
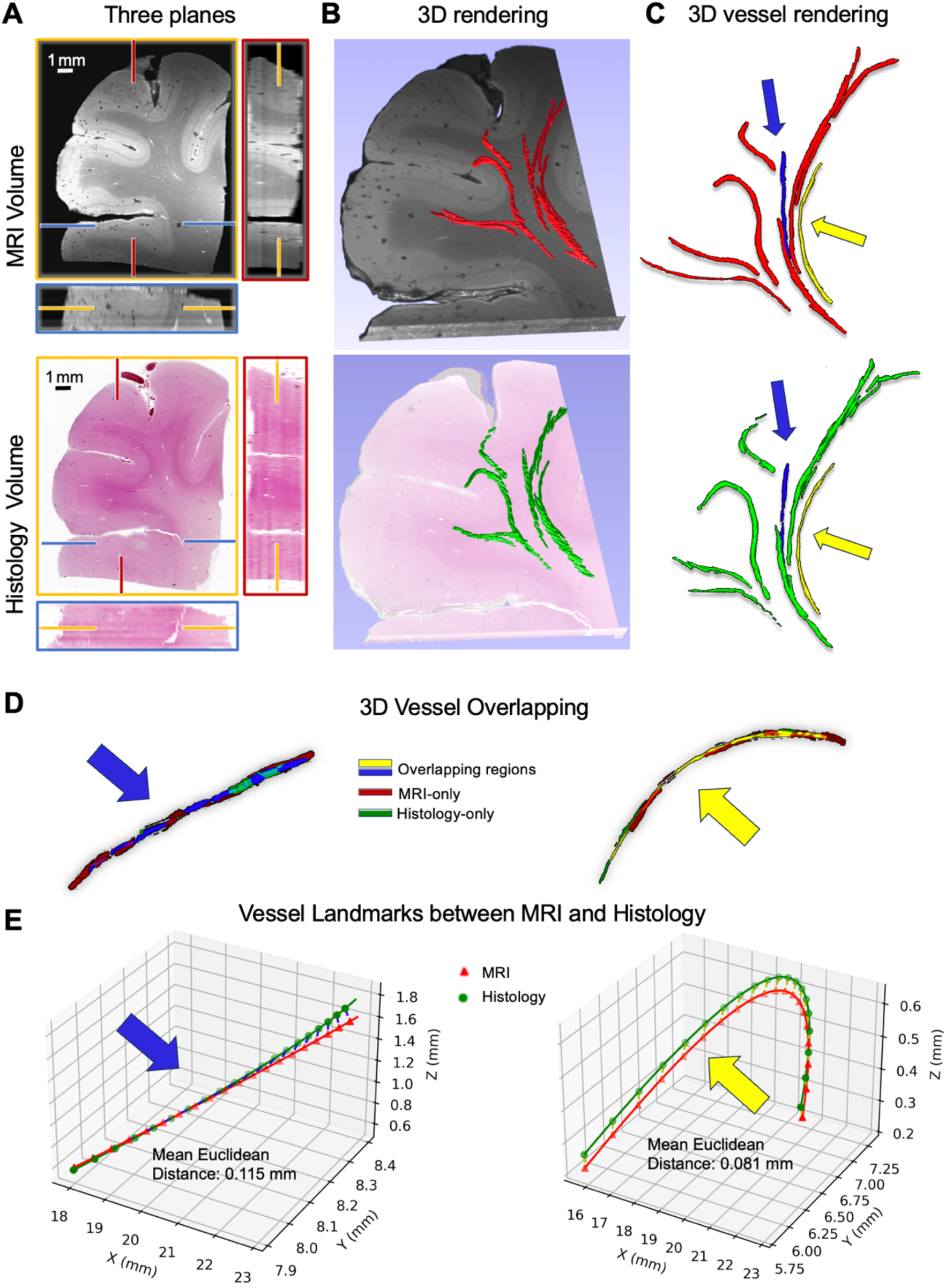
Reconstruction of volumetric histology from Human 2-mm Cortex. **A** shows the reconstructed volumes, while **B** illustrates rendering using 3D Slicer. The manually segmented vessels in both MRI (red) and histology volumes (green) are highlighted in **C** and overlaid on **B**, with two continuous vessels (blue and yellow) selected to assess voxel overlap in **D** and mean Euclidean distances in **E.**

## 4. Discussion

In this work, we presented a novel pipeline, Brewsters Blockface Quantification (BBQ), for high-fidelity alignment between *ex vivo* MRI and histology. Using Brewster’s angle blockface imaging as an intermediate step, we divided the challenging registration problem into more tractable 3D-3D and 2D-2D registration steps. Comparing multiple blockface imaging techniques, we found superior imaging using polarized optics at an angle slightly off Brewster’s angle. TIRL’s Nonlinear 3D and 2D alignment tools delivered high-fidelity 2D registrations of histological details at the level of MRI resolution, shown across an array of samples from two species.

### 4.1. Optimizing Illumination with Brewster’s Adjacent for Enhanced Tissue Contrast

We build upon the off-axis approach (Shojaii et al., 2014) to take advantage of Brewster’s physics to avoid through-plane optical transmission. However, illuminating precisely at Brewster’s angle often results in reduced tissue contrast. This angle is contingent on the refractive index of the materials involved, which is challenging to determine precisely for mixed dehydrated tissues and paraffin in histology blocks. We found that positioning our light source slightly off Brewster’s angle provided superior tissue contrast, likely because it was not fully reflected by the paraffin on the very surface and could reach and be reflected off the tissue too, thus avoiding the dominance of line artifacts on paraffin caused by the blade when illuminating directly at Brewster’s angle. We did observe a relative inversion of gray-white contrast between the exact and slightly off Brewster angle observed in **Fig. 4**. We presume this relates to slight differences in the refractive index of paraffin-embedded white and gray matter, with white matter more closely matching that of paraffin (Biswas & Luu, 2009).

From the perspective of physical apparatus, our protocol setup ensures the acquisition of high-quality sections and images. While our method is limited to specific regions rather than the whole brain, our future work can combine both methods to achieve a comprehensive and accurate MRI-histology pipeline for whole-brain biomarker analysis (Alkemade et al., 2022). An alternative blockface approach includes performing vibratome section while performing polarization-sensitive optical coherence tomography (PS-OCT) (Magnain et al., 2013; Pircher et al., 2011; Wang et al., 2014). OCT can be performed directly on tissue blocks, providing full 3D representations of brain structures, including neurons, fibers, vasculature, and tumors. However, such sectioning setups require the use of a vibratome, which usually generates thicker sections than those used by traditional histology. Devices such as the CoMBI system (Ishii et al., 2021; Tajika et al., 2023) does not use Brewster’s angle physics, so blurring occurs along the z-axis, and there is less room for an operator to collect slides and work with the block. This is particularly important for cutting entire blocks of human brain tissue, which we found requires humidification to maintain section quality. Combining blockface imaging with histological section scans has also been reported in the context of 3D Polarized Light Imaging (3D-PLI), a section-based label-free microscopy technique for studying the brain’s fiber architecture (Ali et al., 2018; Axer et al., 2011; Schmitz et al., 2018; Zeineh et al., 2017). A 3D reconstructed blockface volume provides a perfect reference for the volumetric alignment of brain sections scanned and analyzed with 3D-PLI. However, in contrast to the paraffin-embedded tissue samples used for the pipeline described here, 3D-PLI requires cryo-sectioning and no Brewster setup because of the different reflectivity properties of the paraffin and frozen blocks.

### 4.2. Overcoming Slice-to-Volume Limitations via Blockface Volume Registration

Our proposed computational pipeline achieved robust and precise 2D histology to 3D MRI alignment. Some have used FFD registration in baboons (Dauguet et al., 2007) and mice (Dubois et al., 2007; Lebenberg et al., 2010); we build upon this knowledge by Brewster’s optics and TIRL’s MIND-based registration algorithm. TIRL was first developed to be used to perform deformable 2D-to-3D registration by undistorting and guiding the integration of each histology image into the MRI volume, using two intermediate, standalone brain slab photographs as a bridge, not blockface imaging. However, the accuracy and robustness of TIRL depend significantly on the space searching throughout the MRI space to find a 3D surface that best represents the “cutting plane” for the individual brain slab. This search leverages the unique cross-sectional anatomy of the brain slabs and becomes challenging for small regional slabs, which may have similar cross-sectional anatomical features. Our approach reconstructed serial blockface images into a 3D volume, simplifying the searching problem directly to a 3D-to-3D whole-volume registration process, which ultimately led to higher accuracy required by the precise pathological correlation with MRI biomarkers.

### 4.3. Limitations and Challenges

There are several considerations and challenges for our proposed pipeline. It does require precise positioning of the camera, light source, and microtome, for which we provide detailed guidance (See Supplementary Protocols). TIRL alignment requires parameter optimization (e.g., downsampling) to achieve accurate registration, also included in our scripts. Tissue edges remain a challenge because of often incomplete sampling, but appropriately positioning flat tissue face-down in the embedding mold minimizes this at the front of the block. Thicker tissues may require a deeper 1 cm embedding mold, which we have used successfully (Human 4-mm cortex block). The raw blockface image showed intensity biases (top-bottom gradient) as shown in Supplementary Figures S1 and S2, that nevertheless do not cause a problem for our registration pipelines. We have recently utilized a larger and more uniform light source to improve illumination homogeneity and image quality (See Supplementary Protocols). Our pipeline requires cutting through the entire block and capturing images of each surface slice. While time-consuming, it ensures precise alignment between histology images and volumetric MRI, and paraffin sections can be stored for future use. A single trained individual can perform one blockface sectioning run in one working day. Synchronization of photography with automatic microtomes may further streamline the workflow, with the limiting step being slide capture. The typical practice of placing blocks on ice before sectioning is not practical for consecutive whole-block sectioning; instead, a cooling block and humidification-maintained section quality. Larger tissue blocks may require a larger homogeneous light source illuminating at Brewster’s Adjacent angle. While we do not acquire orientation information on individual slices as in PS-OCT, slides can be subsequently analyzed with Computational Scattered Light Imaging (ComSLI) (Georgiadis et al., 2024; Menzel et al., 2021) or Structure Tensor Analysis (STA) (Budde & Frank, 2012; Schurr & Mezer, 2021). Frozen sectioning is typically performed in OCT, which is white and opaque, so *en face* blockface imaging (**Supplement Fig. S1** *en face* white paraffin) may be more suitable than adapting the optics to a cryotome. H&E staining offers limited gray/white matter contrast, which restricts the effectiveness of conventional registration methods such as Mutual Information-based SyN that primarily rely on outer tissue boundaries. In contrast, the use of the MIND descriptor in TIRL facilitated alignment of multiple stains to MRI, accounting for differences in staining contrast, even when overall contrast was low (**Supplement Fig. S5**).

### 4.4. Future Work

This pipeline is immediately applicable to formalin-fixed, paraffin-embedded brain tissue from any species. We will extend our prior work (Tran et al., 2022; Zeineh et al., 2015) to assess iron mapping in combination with histology markers across hippocampal subfields in AD, which could facilitate ultra-high resolution *in vivo* visualization of iron biomarkers (DiGiacomo et al., 2020). Similar applications exist for motor neuron disorders, e.g., ALS (Kwan et al., 2012) and iron or neuromelanin in the brainstem in PD (Blazejewska et al., 2013; Brammerloh et al., 2022; Lee et al., 2016). The same methodology could be applied to other disorders, including the investigation of animal models of traumatic brain injury: this will provide the histological gold standard to reference *ex vivo* imaging, which should inform feasibility and resolution requirements for *in vivo* imaging. To evaluate the generalizability of our registration pipeline across different histological stains, we tested adjacent sections stained with Hematoxylin, Luxol Fast Blue (LFB), Nissl, Iron (Perls), and Iron (Perls with DAB) which were co-registered with MRI (See **Supplement Fig. S5**). Despite the varying contrasts and structural features of these stains, each stain was aligned to the corresponding MRI slice with high anatomical fidelity, further supporting our pipeline’s applicability for multimodal neuropathological investigations beyond H&E. This approach can also facilitate both MRI segmentation and histological delineation of brain tissue (Casamitjana et al., 2024; Wagstyl et al., n.d.; Wuestefeld et al., 2024). In the future, artificial intelligence could be integrated to enhance registration accuracy and automate segmentation for more efficient and precise quantification.

## 5. Conclusion

Our integrated pipeline BBQ has the potential to provide sub-millimeter, close to micrometer-level registration between MR and densely sampled histology images. The utilization of polarization and Brewster’s optics in blockface imaging significantly enhances contrast and reduces obfuscation of the blockface volume by out-of-plane tissue. TIRL’s MIND-based coregistration is synergistic with our high-quality histology acquisition protocol. The robust coregistration with thorough evaluation provides a reliable framework for correlating histological findings with MRI-detected anomalies in Alzheimer’s disease and beyond.

## Supporting information

Supplementary Protocols

## Data and Code Availability

The raw MRI, Blockface, and Histology data of the Human 2-mm Cortex are all available in the data repository Dryad: http://datadryad.org/share/r5MCv5UGxOd5B11V22NPRVOYuzy4vLlH1pMR-IvgswU. Other specimen data are available from the corresponding author by request. All codes used for the processing and registration in this study are provided via Gitlab (https://code.stanford.edu/zeinehlab/BBQ.git). The authors declare that they have no known competing financial interests.

## Author Contributions

M.Z., M.G., Y.W., P.D., and P.S. were responsible for the conception and design of the study. M.G., Y.W., W.H., I.N.H., P.D., H.M.T., L.T., M.C., N.N., S.L., D.B.C., J.N., and M.Z. were responsible for data curation and analysis. Y.W., I.N.H., W.S., M.R., and M.Z. were responsible for software development. Y.W., L.T., N.N., S.L., and M.Z. contributed to data segmentation and evaluation. Y.W., W.H., I.N.H., P.D., H.M.T., M.C., P.S., M.A., W.S., M.R., I.C., J.N., M.G., and M.Z. contributed to writing, reviewing, and editing. M.G. co-supervised the study. M.Z. supervised the study and acquired funding.

## Acknowledgments

The authors thank Christina Tisca and Aurea B. Martins-Bach for their valuable advice on tissue optimization. The present work was supported by the National Institutes of Health (NIH), award numbers R01AG061120-01 (M.Z.) and R03AG083702 (M.G.), Alzheimer’s Association Research Fellowship no. 24AARF-1241479 (M.G), Taube Stanford Children’s Concussion Initiative (M.Z.), Wu Tsai Human Performance Agility Project Program and PHIND Pilot Program (M.Z.).

## Supplementary Materials

**Supplement Figure S1:**
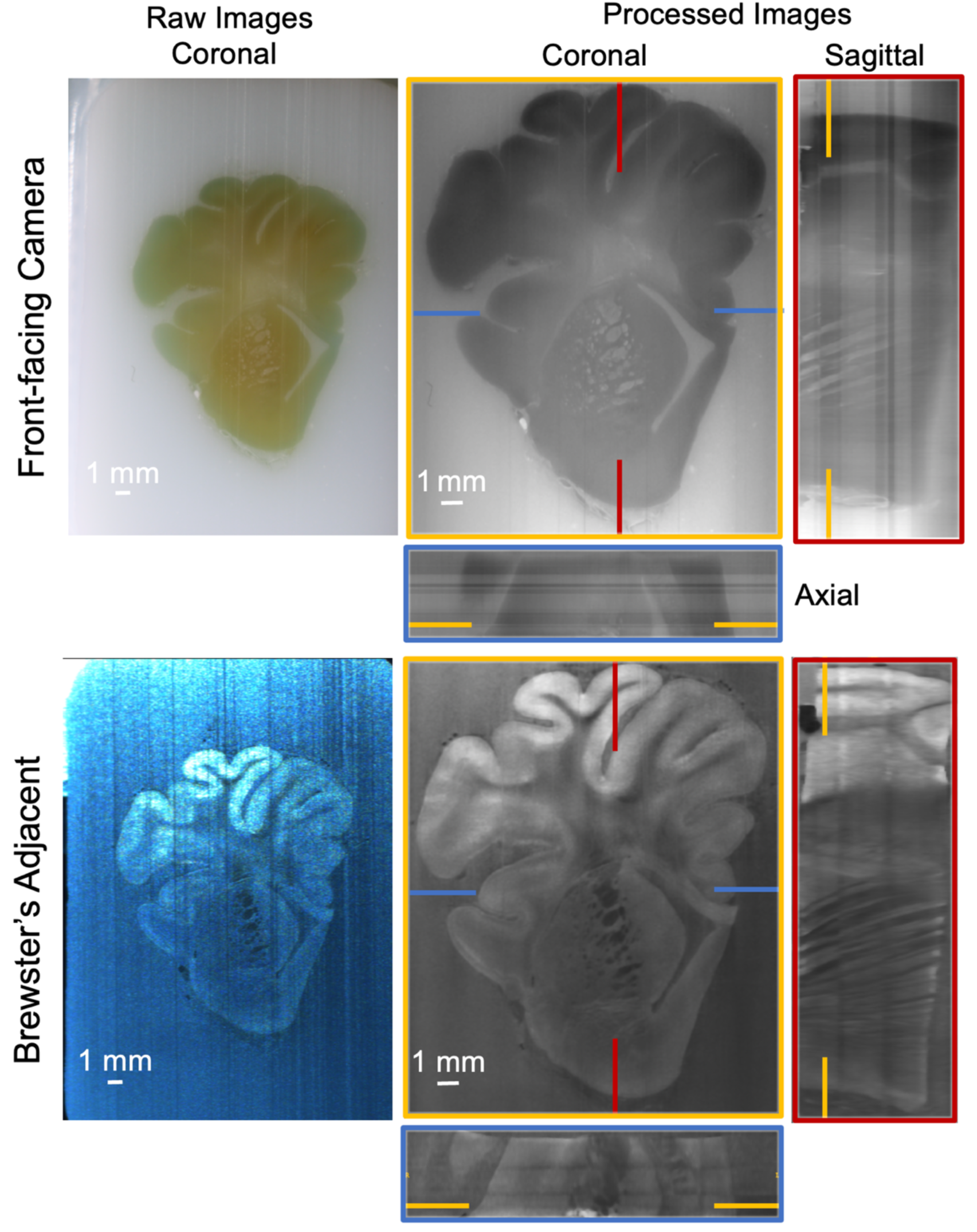
Comparison of raw and processed blockface images of the white-paraffin embedded pig coronal slab captured using two different lighting setups, both designed to eliminate depth information: a front-facing camera and a Brewster’s angle adjacent camera.

**Supplement Figure S2:**
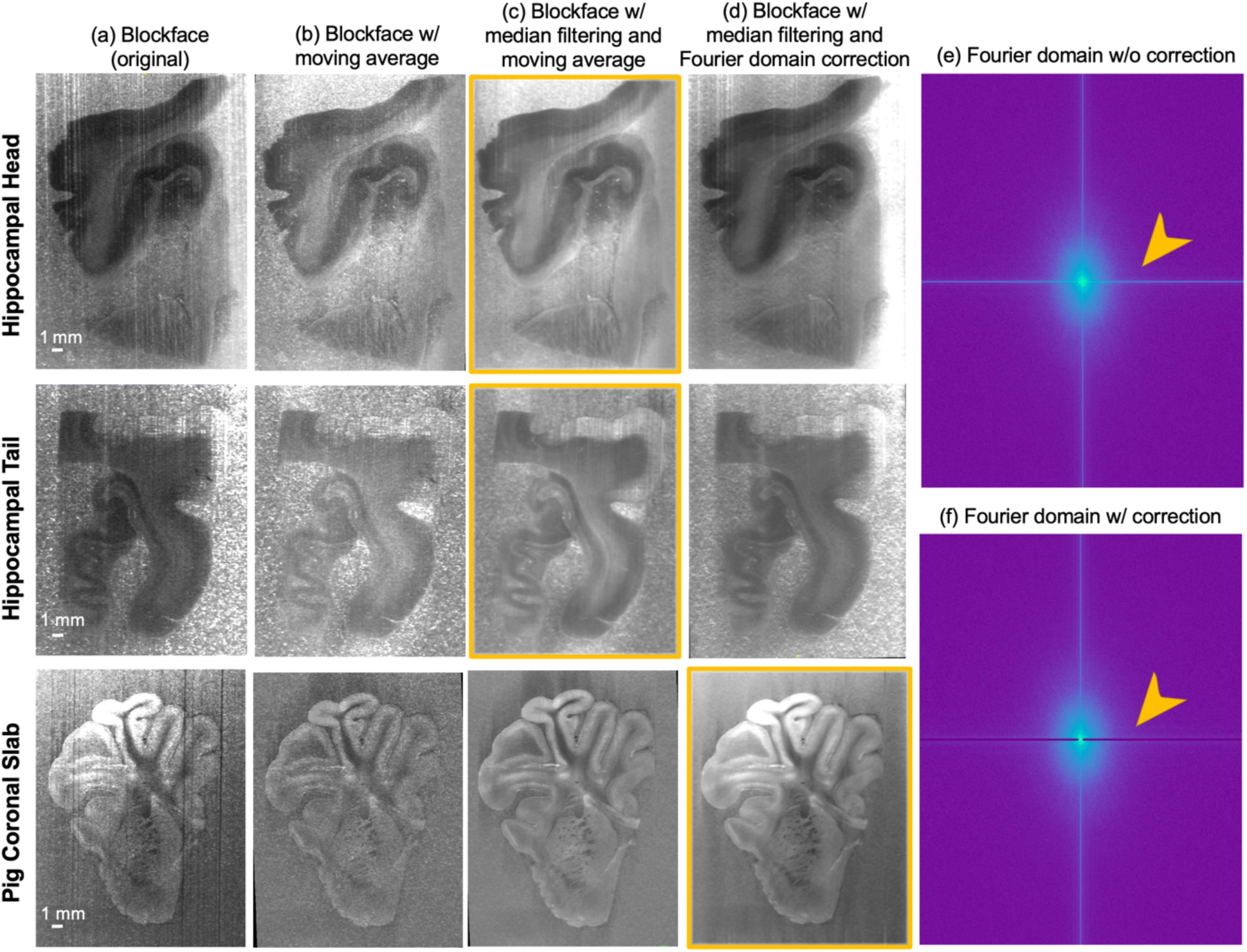
Blockface Imaging Processing. (a) Reconstructed and cropped blockface volumes. To address the vertical line artifact from sectioning, (b) a moving average method was applied in the spatial domain (c) with subsequent median filtering, or (d) a Fourier domain correction was adopted, where signals of the line artifacts were filtered out (pointed by orange arrows) in the Fourier domain (e, f). Images within the orange frames were identified as the best quality and selected for subsequent registration with MRI.

**Supplement Figure S3:**
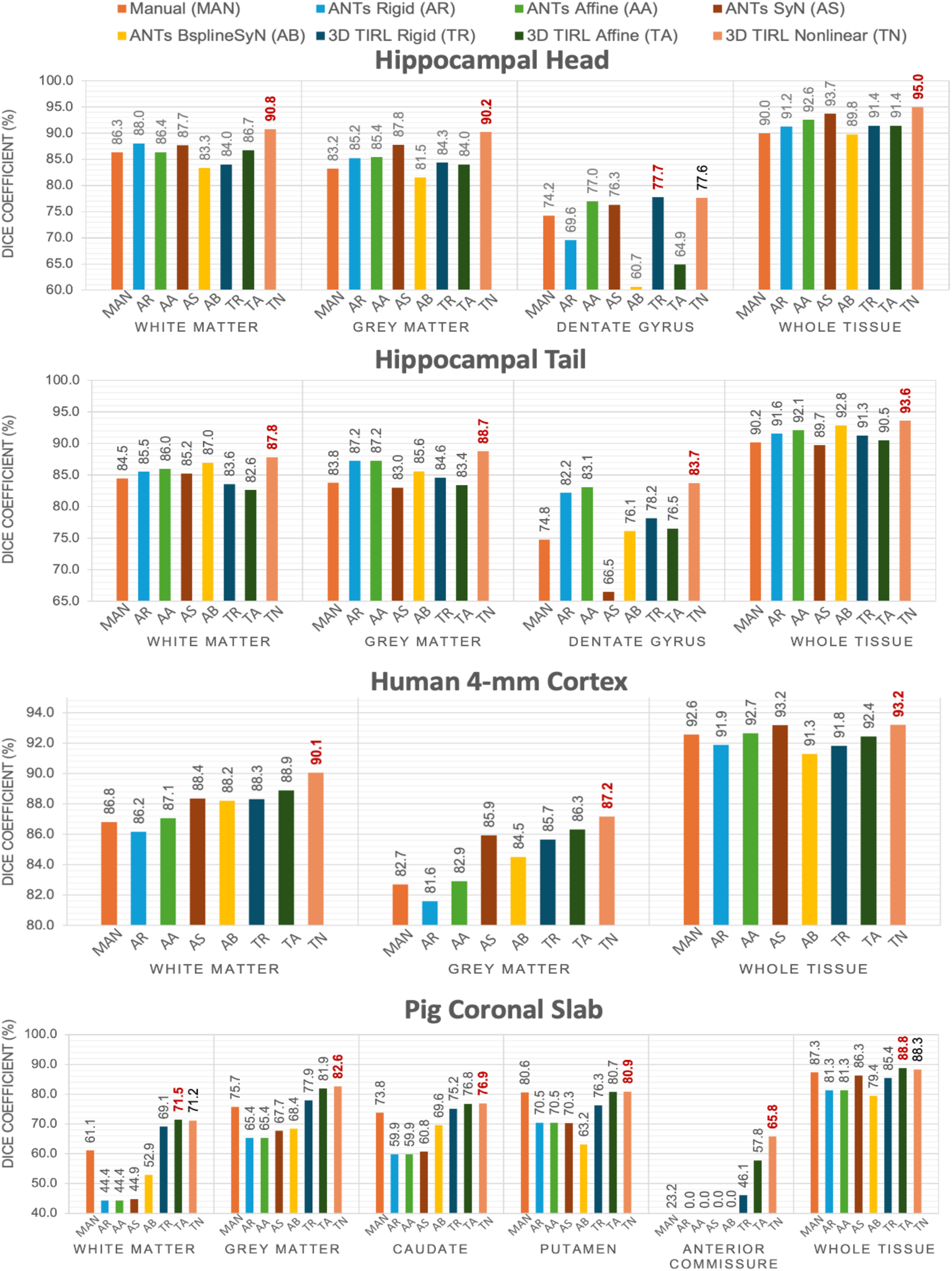
Dice similarity coefficient scores (%) of the segmentations of MRI and blockface volumes after coregistration across different specimens. “Whole Tissue” represents the score for the entire composite area. The highest score (best overlap) is indicated in Red.

**Supplement Figure S4:**
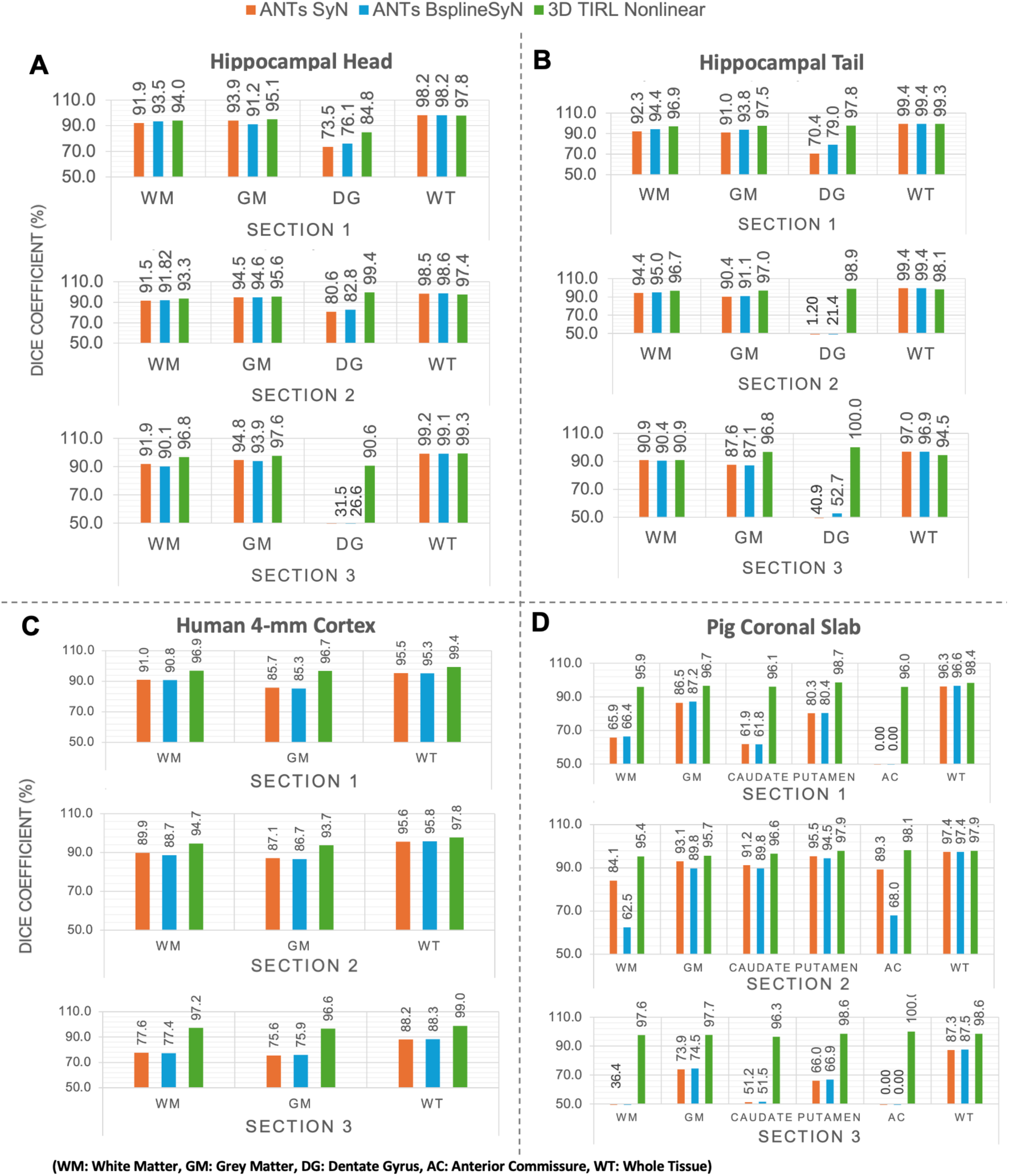
Dice similarity coefficient (%) analysis for 2D registration of MRI and histology across multiple hippocampal head, tail, 4-mm cortex and pig coronal slab slides with H&E staining. TIRL shows a superior performance in the dentate gyrus (DG) and Anterior Commissure (AC).

**Supplement Figure S5:**
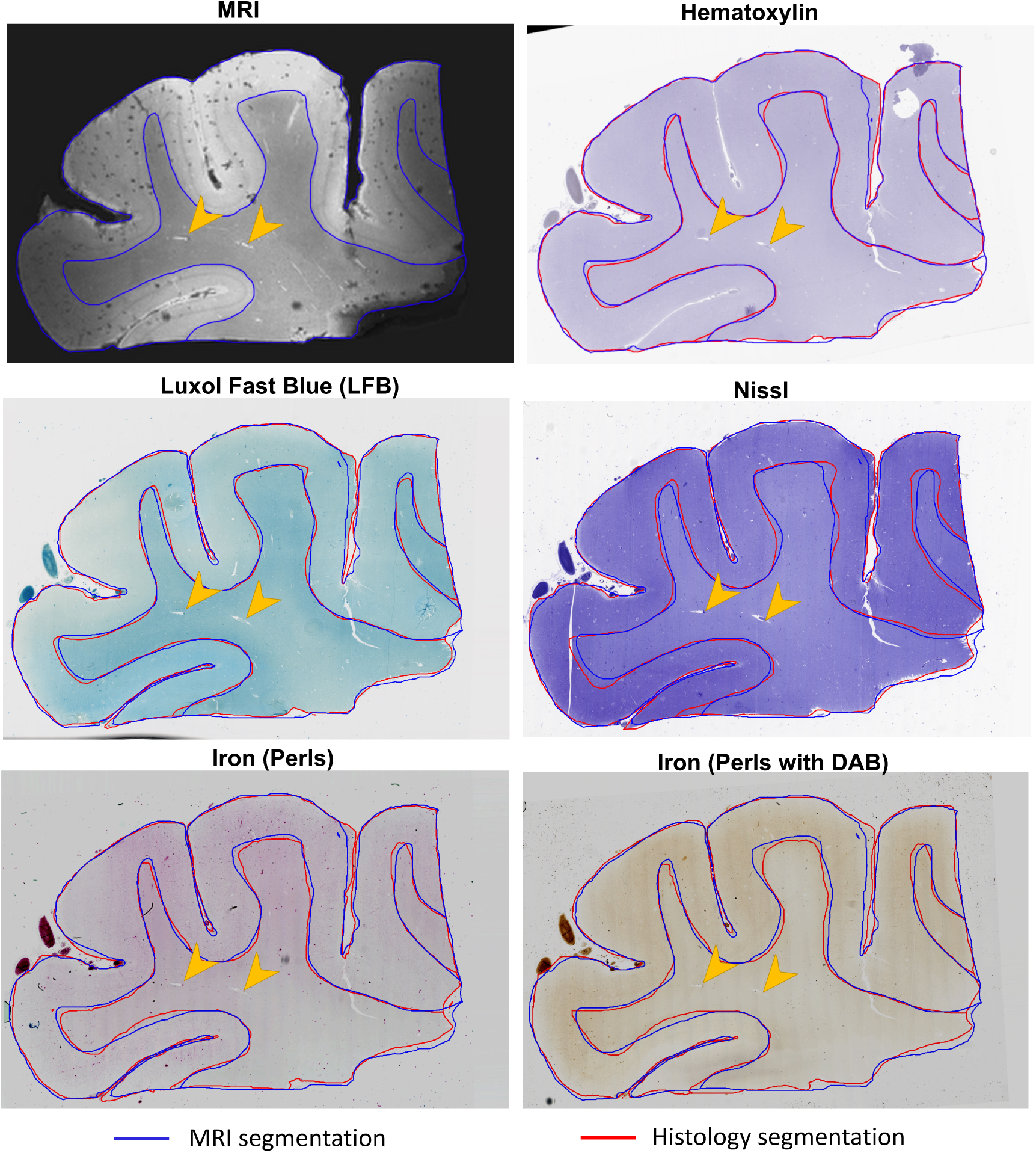
Adjacent sections from Human 2-mm Cortex stained with Hematoxylin, Luxol Fast Blue (LFB), Nissl, Iron (Perls), and Iron (Perls with DAB) were shown registered to MRI (top left). White matter and grey matter were manually segmented on MRI slices and histology sections transformed into MRI space. Vessels were marked by orange arrows as internal landmarks. Brightness and contrast were adjusted for Iron (Perls) and Iron (Perls with DAB) to enhance stain visibility in the figure.

## Footnotes

1 https://www.mrtrix.org/

## REFERENCES

1. Adler, D. H., Pluta, J., Kadivar, S., Craige, C., Gee, J. C., Avants, B. B., & Yushkevich, P. A. (2014). Histology-derived volumetric annotation of the human hippocampal subfields in postmortem MRI. NeuroImage, 84, 505–523. 10.1016/j.neuroimage.2013.08.067

2. Ali, S., Wörz, S., Amunts, K., Eils, R., Axer, M., & Rohr, K. (2018). Rigid and non-rigid registration of polarized light imaging data for 3D reconstruction of the temporal lobe of the human brain at micrometer resolution. NeuroImage, 181, 235–251. 10.1016/j.neuroimage.2018.06.084

3. Alkemade, A., Bazin, P.-L., Balesar, R., Pine, K., Kirilina, E., Möller, H. E., Trampel, R., Kros, J. M., Keuken, M. C., Bleys, R. L. A. W., Swaab, D. F., Herrler, A., Weiskopf, N., & Forstmann, B. U. (2022). A unified 3D map of microscopic architecture and MRI of the human brain. Science Advances. 10.1126/sciadv.abj7892

4. Amunts, K., Kedo, O., Kindler, M., Pieperhoff, P., Mohlberg, H., Shah, N. J., Habel, U., Schneider, F., & Zilles, K. (2005). Cytoarchitectonic mapping of the human amygdala, hippocampal region and entorhinal cortex: Intersubject variability and probability maps. Anatomy and Embryology, 210(5), 343–352. 10.1007/s00429-005-0025-5

5. Avants, B., Tustison, N. J., & Song, G. (2009). Advanced Normalization Tools: V1.0. The Insight Journal. 10.54294/uvnhin

6. Axer, M., Amunts, K., Grässel, D., Palm, C., Dammers, J., Axer, H., Pietrzyk, U., & Zilles, K. (2011). A novel approach to the human connectome: Ultra-high resolution mapping of fiber tracts in the brain. NeuroImage, 54(2), 1091–1101. 10.1016/j.neuroimage.2010.08.075

7. Barthel, H., Schroeter, M. L., Hoffmann, K.-T., & Sabri, O. (2015). PET/MR in dementia and other neurodegenerative diseases. Seminars in Nuclear Medicine, 45(3), 224–233. 10.1053/j.semnuclmed.2014.12.003

8. Biswas, T. K., & Luu, T. M. (2009). In vivo MR Measurement of Refractive Index, Relative Water Content and T2 Relaxation time of Various Brain lesions With Clinical Application to Discriminate Brain Lesions. The Internet Journal of Radiology, 13(1). https://ispub.com/IJRA/13/1/8277

9. Blazejewska, A. I., Schwarz, S. T., Pitiot, A., Stephenson, M. C., Lowe, J., Bajaj, N., Bowtell, R. W., Auer, D. P., & Gowland, P. A. (2013). Visualization of nigrosome 1 and its loss in PD: Pathoanatomical correlation and in vivo 7 T MRI. Neurology, 81(6), 534–540. 10.1212/WNL.0b013e31829e6fd2

10. Bobinski, M., de Leon, M. J., Wegiel, J., Desanti, S., Convit, A., Saint Louis, L. A., Rusinek, H., & Wisniewski, H. M. (2000). The histological validation of post mortem magnetic resonance imaging-determined hippocampal volume in Alzheimer’s disease. Neuroscience, 95(3), 721–725. 10.1016/s0306-4522(99)00476-5

11. Brammerloh, M., Kirilina, E., Alkemade, A., Bazin, P.-L., Jantzen, C., Jäger, C., Herrler, A., Pine, K. J., Gowland, P. A., Morawski, M., Forstmann, B. U., & Weiskopf, N. (2022). Swallow Tail Sign: Revisited. Radiology, 305(3), 674–677. 10.1148/radiol.212696

12. Brewster, D. (1815). On the Laws Which Regulate the Polarisation of Light by Reflexion from Transparent Bodies. Philosophical Transactions of the Royal Society of London, 105, 125–159.

13. Budde, M. D., & Frank, J. A. (2012). Examining brain microstructure using structure tensor analysis of histological sections. NeuroImage, 63(1), 1–10. 10.1016/j.neuroimage.2012.06.042

14. Casamitjana, A., Mancini, M., Robinson, E., Peter, L., Annunziata, R., Althonayan, J., Crampsie, S., Blackburn, E., Billot, B., Atzeni, A., Puonti, O., Balbastre, Y., Schmidt, P., Hughes, J., Augustinack, J. C., Edlow, B. L., Zöllei, L., Thomas, D. L., Kliemann, D., … Iglesias, J. E. (2024). A next-generation, histological atlas of the human brain and its application to automated brain MRI segmentation (p. 2024.02.05.579016). bioRxiv. 10.1101/2024.02.05.579016

15. Casero, R., Siedlecka, U., Jones, E. S., Gruscheski, L., Gibb, M., Schneider, J. E., Kohl, P., & Grau, V. (2017). Transformation diffusion reconstruction of three-dimensional histology volumes from two-dimensional image stacks. Medical Image Analysis, 38, 184–204. 10.1016/j.media.2017.03.004

16. Chakravarty, M. M., Bertrand, G., Hodge, C. P., Sadikot, A. F., & Collins, D. L. (2006). The creation of a brain atlas for image guided neurosurgery using serial histological data. NeuroImage, 30(2), 359–376. 10.1016/j.neu-roimage.2005.09.041

17. Cosottini, M., Donatelli, G., Costagli, M., Caldarazzo Ienco, E., Frosini, D., Pesaresi, I., Biagi, L., Siciliano, G., & Tosetti, M. (2016). High-Resolution 7T MR Imaging of the Motor Cortex in Amyotrophic Lateral Sclerosis. AJNR: American Journal of Neuroradiology, 37(3), 455–461. 10.3174/ajnr.A4562

18. Dauguet, J., Delzescaux, T., Condé, F., Mangin, J.-F., Ayache, N., Hantraye, P., & Frouin, V. (2007). Three-dimensional reconstruction of stained histological slices and 3D non-linear registration with *in-vivo* MRI for whole baboon brain. Journal of Neuroscience Methods, 164(1), 191–204. 10.1016/j.jneumeth.2007.04.017

19. De Barros, A., Arribarat, G., Combis, J., Chaynes, P., & Péran, P. (2019). Matching ex vivo MRI With Iron Histology: Pearls and Pitfalls. Frontiers in Neuroanatomy, 13. 10.3389/fnana.2019.00068

20. Dice, L. R. (1945). Measures of the Amount of Ecologic Association Between Species. Ecology, 26(3), 297–302. 10.2307/1932409

21. DiGiacomo, P., Maclaren, J., Aksoy, M., Tong, E., Carlson, M., Lanzman, B., Hashmi, S., Watkins, R., Rosenberg, J., Burns, B., Skloss, T. W., Rettmann, D., Rutt, B., Bammer, R., & Zeineh, M. (2020). A within-coil optical prospective motion-correction system for brain imaging at 7T. Magnetic Resonance in Medicine, 84(3), 1661–1671. 10.1002/mrm.28211

22. Dubois, A., Dauguet, J., Herard, A.-S., Besret, L., Duchesnay, E., Frouin, V., Hantraye, P., Bonvento, G., & Delzescaux, T. (2007). Automated three-dimensional analysis of histological and autoradiographic rat brain sections: Application to an activation study. Journal of Cerebral Blood Flow and Metabolism: Official Journal of the International Society of Cerebral Blood Flow and Metabolism, 27(10), 1742– 1755. 10.1038/sj.jcbfm.9600470

23. Fedorov, A., Beichel, R., Kalpathy-Cramer, J., Finet, J., Fillion-Robin, J.-C., Pujol, S., Bauer, C., Jennings, D., Fennessy, F., Sonka, M., Buatti, J., Aylward, S., Miller, J. V., Pieper, S., & Kikinis, R. (2012). 3D Slicer as an image computing platform for the Quantitative Imaging Network. Magnetic Resonance Imaging, 30(9), 1323–1341. 10.1016/j.mri.2012.05.001

24. Georgiadis, M., Auf der Heiden, F., Abbasi, H., Ettema, L., Nirschl, J., Moein Taghavi, H., Wakatsuki, M., Liu, A., Ho, W. H. D., Carlson, M., Doukas, M., Koppes, S. A., Keereweer, S., Sobel, R. A., Setsompop, K., Liao, C., Amunts, K., Axer, M., Ze-ineh, M., & Menzel, M. (2024). Micron-resolution fiber mapping in histology independent of sample preparation. bioRxiv: The Preprint Server for Biology, 2024.03.26.586745. 10.1101/2024.03.26.586745

25. Gibson, E., Crukley, C., Gaed, M., Gómez, J. A., Moussa, M., Chin, J. L., Bauman, G. S., Fenster, A., & Ward, A. D. (2012). Registration of prostate histology images to ex vivo MR images via strand-shaped fiducials. Journal of Magnetic Resonance Imaging, 36(6), 1402–1412. 10.1002/jmri.23767

26. Goubran, M., Crukley, C., de Ribaupierre, S., Peters, T. M., & Khan, A. R. (2013). Image registration of ex-vivo MRI to sparsely sectioned histology of hippocampal and neocortical temporal lobe specimens. NeuroImage, 83, 770–781. 10.1016/j.neuroimage.2013.07.053

27. Groen, H. C., van Walsum, T., Rozie, S., Klein, S., van Gaalen, K., Gijsen, F. J. H., Wielopolski, P. A., van Beusekom, H. M. M., de Crom, R., Verhagen, H. J. M., van der Steen, A. F. W., van der Lugt, A., Wentzel, J. J., & Niessen, W. J. (2010). Three-dimensional registration of histology of human atherosclerotic carotid plaques to *in-vivo* imaging. Journal of Biomechanics, 43(11), 2087–2092. 10.1016/j.jbiomech.2010.04.005

28. Heinrich, M. P., Jenkinson, M., Bhushan, M., Matin, T., Gleeson, F. V., Brady, S. M., & Schnabel, J. A. (2012). MIND: Modality independent neighbourhood descriptor for multi-modal deformable registration. Medical Image Analysis, 16(7), 1423– 1435. 10.1016/j.media.2012.05.008

29. Hnilicova, P., Kantorova, E., Sutovsky, S., Grofik, M., Zelenak, K., Kurca, E., Zilka, N., Parvanovova, P., & Kolisek, M. (2023). Imaging Methods Applicable in the Diagnostics of Alzheimer’s Disease, Considering the Involvement of Insulin Resistance. International Journal of Molecular Sciences, 24(4), 3325. 10.3390/ijms24043325

30. Howard, A. F. D., Huszar, I. N., Smart, A., Cottaar, M., Daubney, G., Hanayik, T., Khrapitchev, A. A., Mars, R. B., Mollink, J., Scott, C., Sibson, N. R., Sallet, J., Jbabdi, S., & Miller, K. L. (2023). An open resource combining multi-contrast MRI and microscopy in the macaque brain. Nature Communications, 14(1), 4320. 10.1038/s41467-023-39916-1

31. Hui, R. (2020). Chapter 2—Optical fibers. In R. Hui (Ed.), Introduction to Fiber-Optic Communications (pp. 19–76). Academic Press. 10.1016/B978-0-12-805345-4.00002-0

32. Huszar, I. N., Pallebage-Gamarallage, M., Bangerter-Christensen, S., Brooks, H., Fitz-gibbon, S., Foxley, S., Hiemstra, M., Howard, A. F. D., Jbabdi, S., Kor, D. Z. L., Leonte, A., Mollink, J., Smart, A., Tendler, B. C., Turner, M. R., Ansorge, O., Miller, K. L., & Jenkinson, M. (2023). Tensor image registration library: Deformable registration of stand-alone histology images to whole-brain post-mortem MRI data. NeuroImage, 265, 119792. 10.1016/j.neu-roimage.2022.119792

33. Iglesias, J. E., Modat, M., Peter, L., Stevens, A., Annunziata, R., Vercauteren, T., Lein, E., Fischl, B., Ourselin, S., & Alzheimer’s Disease Neuroimaging Initiative. (2018). Joint registration and synthesis using a probabilistic model for alignment of MRI and histological sections. Medical Image Analysis, 50, 127–144. 10.1016/j.media.2018.09.002

34. Ishii, N., Tajika, Y., Murakami, T., Galipon, J., Shirahata, H., Mukai, R., Uehara, D., Kaneko, R., Yamazaki, Y., Yoshimoto, Y., & Iwasaki, H. (2021). Correlative microscopy and block-face imaging (CoMBI) method for both paraffin-embedded and frozen specimens. Scientific Reports, 11(1), 13108. 10.1038/s41598-021-92485-5

35. Johnson, G. A., Badea, A., Brandenburg, J., Cofer, G., Fubara, B., Liu, S., & Nissanov, J. (2010). Waxholm Space: An image-based reference for coordinating mouse brain research. NeuroImage, 53(2), 365–372. 10.1016/j.neu-roimage.2010.06.067

36. Kellner, E., Dhital, B., Kiselev, V. G., & Reisert, M. (2016). Gibbs-ringing artifact removal based on local subvoxel-shifts. Magnetic Resonance in Medicine, 76(5), 1574– 1581. 10.1002/mrm.26054

37. Kim, T.-S., Singh, M., Sungkarat, W., Zarow, C., & Chui, H. (2000). Automatic registration of postmortem brain slices to MRI reference volume. IEEE Transactions on Nuclear Science, 47(4), 1607–1613. IEEE Transactions on Nuclear Science. 10.1109/23.873023

38. Kwan, J. Y., Jeong, S. Y., Van Gelderen, P., Deng, H.-X., Quezado, M. M., Danielian, L. E., Butman, J. A., Chen, L., Bayat, E., Russell, J., Siddique, T., Duyn, J. H., Rouault, T. A., & Floeter, M. K. (2012). Iron accumulation in deep cortical layers accounts for MRI signal abnormalities in ALS: Correlating 7 tesla MRI and pathol-ogy. PloS One, 7(4), e35241. 10.1371/journal.pone.0035241

39. Lebenberg, J., Hérard, A.-S., Dubois, A., Dauguet, J., Frouin, V., Dhenain, M., Hantraye, P., & Delzescaux, T. (2010). Validation of MRI-based 3D digital atlas registration with histological and autoradiographic volumes: An anatomofunctional transgenic mouse brain imaging study. NeuroImage, 51(3), 1037–1046. 10.1016/j.neuroimage.2010.03.014

40. Lee, J.-H., Baek, S.-Y., Song, Y., Lim, S., Lee, H., Nguyen, M. P., Kim, E.-J., Huh, G. Y., Chun, S. Y., & Cho, H. (2016). The Neuromelanin-related T2* Contrast in Postmortem Human Substantia Nigra with 7T MRI. Scientific Reports, 6, 32647. 10.1038/srep32647

41. Magnain, C., Augustinack, J. C., Reuter, M., Wachinger, C., Frosch, M. P., Ragan, T., Akkin, T., Wedeen, V. J., Boas, D. A., & Fischl, B. (2013). Blockface Histology with Optical Coherence Tomography: A Comparison with Nissl Staining. NeuroImage, 84, 524. 10.1016/j.neuroimage.2013.08.072

42. Mancini, M., Casamitjana, A., Peter, L., Robinson, E., Crampsie, S., Thomas, D. L., Holton, J. L., Jaunmuktane, Z., & Iglesias, J. E. (2020). A multimodal computational pipeline for 3D histology of the human brain. Scientific Reports, 10(1), 13839. 10.1038/s41598-020-69163-z

43. Menzel, M., Reuter, J. A., Gräßel, D., Huwer, M., Schlömer, P., Amunts, K., & Axer, M. (2021). Scattered Light Imaging: Resolving the substructure of nerve fiber crossings in whole brain sections with micrometer resolution. NeuroImage, 233, 117952. 10.1016/j.neuroimage.2021.117952

44. Nir, G., Sahebjavaher, R. S., Kozlowski, P., Chang, S. D., Jones, E. C., Goldenberg, S. L., & Salcudean, S. E. (2014). Registration of Whole-Mount Histology and Volumetric Imaging of the Prostate Using Particle Filtering. IEEE Transactions on Medical Imaging, 33(8), 1601–1613. IEEE Transactions on Medical Imaging. 10.1109/TMI.2014.2319231

45. Ohnishi, T., Nakamura, Y., Tanaka, T., Tanaka, T., Hashimoto, N., Haneishi, H., Batchelor, T. T., Gerstner, E. R., Taylor, J. W., Snuderl, M., & Yagi, Y. (2016). Deformable image registration between pathological images and MR image via an optical macro image. Pathology - Research and Practice, 212(10), 927–936. 10.1016/j.prp.2016.07.018

46. Osechinskiy, S., & Kruggel, F. (2011). Slice-to-Volume Nonrigid Registration of Histological Sections to MR Images of the Human Brain. Anatomy Research International, 2011(1), 287860. 10.1155/2011/287860

47. Pichat, J., Iglesias, J. E., Yousry, T., Ourselin, S., & Modat, M. (2018). A Survey of Methods for 3D Histology Reconstruction. Medical Image Analysis, 46, 73–105. 10.1016/j.media.2018.02.004

48. Pircher, M., Hitzenberger, C. K., & Schmidt-Erfurth, U. (2011). Polarization sensitive optical coherence tomography in the human eye. Progress in Retinal and Eye Research, 30(6), 431. 10.1016/j.preteyeres.2011.06.003

49. Rueckert, D., Sonoda, L. I., Hayes, C., Hill, D. L. G., Leach, M. O., & Hawkes, D. J. (1999). Nonrigid registration using free-form deformations: Application to breast MR images. IEEE Transactions on Medical Imaging, 18(8), 712–721. 10.1109/42.796284

50. Rusu, M., Golden, T., Wang, H., Gow, A., & Madabhushi, A. (2015). Framework for 3D histologic reconstruction and fusion with in vivo MRI: Preliminary results of characterizing pulmonary inflammation in a mouse model. Medical Physics, 42(8), 4822–4832. 10.1118/1.4923161

51. Schilling, K. G., Gao, Y., Stepniewska, I., Janve, V., Landman, B. A., & Anderson, A. W. (2019a). Anatomical accuracy of standard-practice tractography algorithms in the motor system—A histological validation in the squirrel monkey brain. Magnetic Resonance Imaging, 55, 7–25. 10.1016/j.mri.2018.09.004

52. Schilling, K. G., Gao, Y., Stepniewska, I., Janve, V., Landman, B. A., & Anderson, A. W. (2019b). Histologically derived fiber response functions for diffusion MRI vary across white matter fibers—An ex vivo validation study in the squirrel monkey brain. NMR in Biomedicine, 32(6), e4090. 10.1002/nbm.4090

53. Schmitz, D., Amunts, K., Lippert, T., & Axer, M. (2018). A least squares approach for the reconstruction of nerve fiber orientations from tiltable specimen experiments in 3D-PLI. 2018 IEEE 15th International Symposium on Biomedical Imaging (ISBI 2018), 132–135. 10.1109/ISBI.2018.8363539

54. Schormann, T., Dabringhaus, A., & Zilles, K. (1995). Statistics of deformations in histology and application to improved alignment with MRI. IEEE Transactions on Medical Imaging, 14(1), 25–35. IEEE Transactions on Medical Imaging. 10.1109/42.370399

55. Schurr, R., & Mezer, A. A. (2021). The glial framework reveals white matter fiber architecture in human and primate brains. Science, 374(6568), 762–767. 10.1126/science.abj7960

56. Shojaii, R., Bacopulos, S., Yang, W., Karavardanyan, T., Spyropoulos, D., Raouf, A., Martel, A., & Seth, A. (2014). Reconstruction of 3-Dimensional Histology Volume and its Application to Study Mouse Mammary Glands. Journal of Visualized Experiments : JoVE, 89, 51325. 10.3791/51325

57. Smart, A., Tisca, C., Huszar, I. N., Kor, D., Ansorge, O., Tachrount, M., Smart, S., Lerch, J. P., Miller, K. L., & Martins-Bach, A. B. (2023). Protocol for tissue processing and paraffin embedding of mouse brains following *ex vivo* MRI. STAR Protocols, 4(4), 102681. 10.1016/j.xpro.2023.102681

58. Tajika, Y., Ishii, N., Morimura, Y., Fukuda, K., Shikada, M., Murakami, T., Ichinose, S., Yoshimoto, Y., & Iwasaki, H. (2023). Correlative microscopy and block-face imaging (CoMBI): A 3D imaging method with wide applicability in the field of biological science. Anatomical Science International, 98(3), 353–359. 10.1007/s12565-023-00705-x

59. Tran, D., DiGiacomo, P., Born, D. E., Georgiadis, M., & Zeineh, M. (2022). Iron and Alzheimer’s Disease: From Pathology to Imaging. Frontiers in Human Neuroscience, 16, 838692. 10.3389/fnhum.2022.838692

60. Uberti, M., Liu, Y., Dou, H., Mosley, R. L., Gendelman, H. E., & Boska, M. (2009). Registration of in vivo MR to histology of rodent brains using blockface imaging (X. P. Hu & A. V. Clough, Eds.; p. 726213). 10.1117/12.812711

61. Veraart, J., Novikov, D. S., Christiaens, D., Ades-aron, B., Sijbers, J., & Fieremans, E. (2016). Denoising of diffusion MRI using random matrix theory. NeuroImage, 142, 394–406. 10.1016/j.neuroimage.2016.08.016

62. Wagstyl, K., Lepage, C., Bludau, S., Zilles, K., Fletcher, P. C., Amunts, K., & Evans, A. C. (n.d.). Mapping Cortical Laminar Structure in the 3D BigBrain. Retrieved March 11, 2025, from 10.1093/cercor/bhy074

63. Wang, H., Zhu, J., Reuter, M., Vinke, L. N., Yendiki, A., Boas, D. A., Fischl, B., & Akkin, T. (2014). Cross-validation of serial optical coherence scanning and diffusion tensor imaging: A study on neural fiber maps in human medulla oblongata. NeuroImage, 100, 395–404. 10.1016/j.neuroimage.2014.06.032

64. Wuestefeld, A., Baumeister, H., Adams, J. N., de Flores, R., Hodgetts, C. J., Mazloum-Farzaghi, N., Olsen, R. K., Puliyadi, V., Tran, T. T., Bakker, A., Canada, K. L., Dalton, M. A., Daugherty, A. M., La Joie, R., Wang, L., Bedard, M. L., Buendia, E., Chung, E., Denning, A., … Wisse, L. E. M. (2024). Comparison of histological delineations of medial temporal lobe cortices by four independent neuroanatomy laboratories. Hippocampus, 34(5), 241–260. 10.1002/hipo.23602

65. Yang, Z., Richards, K., Kurniawan, N. D., Petrou, S., & Reutens, D. C. (2012). MRI-guided volume reconstruction of mouse brain from histological sections. Journal of Neuroscience Methods, 211(2), 210–217. 10.1016/j.jneumeth.2012.08.021

66. Yelnik, J., Bardinet, E., Dormont, D., Malandain, G., Ourselin, S., Tandé, D., Karachi, C., Ayache, N., Cornu, P., & Agid, Y. (2007). A three-dimensional, histological and deformable atlas of the human basal ganglia. I. Atlas construction based on immunohistochemical and MRI data. NeuroImage, 34(2), 618–638. 10.1016/j.neuroimage.2006.09.026

67. Yushkevich, P. A., Piven, J., Hazlett, H. C., Smith, R. G., Ho, S., Gee, J. C., & Gerig, G. (2006). User-guided 3D active contour segmentation of anatomical structures: Significantly improved efficiency and reliability. NeuroImage, 31(3), 1116–1128. 10.1016/j.neuroimage.2006.01.015

68. Yushkevich, P. A., Wisse, L., Adler, D., Ittyerah, R., Pluta, J. B., Robinson, J. L., Schuck, T., Trojanowski, J. Q., Grossman, M., Detre, J. A., Elliott, M. A., Toledo, J. B., Liu, W., Pickup, S., Das, S. R., & Wolk, D. A. (2016). A framework for informing segmentation of in vivo MRI with information derived from ex vivo imaging: Application in the medial temporal lobe. 2016 38th Annual International Conference of the IEEE Engineering in Medicine and Biology Society (EMBC), 6014–6017. 10.1109/EMBC.2016.7592099

69. Zeineh, M. M., Chen, Y., Kitzler, H. H., Hammond, R., Vogel, H., & Rutt, B. K. (2015). Activated iron-containing microglia in the human hippocampus identified by magnetic resonance imaging in Alzheimer disease. Neurobiology of Aging, 36(9), 2483–2500. 10.1016/j.neurobiolaging.2015.05.022

70. Zeineh, M. M., Palomero-Gallagher, N., Axer, M., Gräßel, D., Goubran, M., Wree, A., Woods, R., Amunts, K., & Zilles, K. (2017). Direct Visualization and Mapping of the Spatial Course of Fiber Tracts at Microscopic Resolution in the Human Hippocampus. Cerebral Cortex (New York, N.Y.: 1991), 27(3), 1779–1794. 10.1093/cercor/bhw010

71. Zhang, X. Y., Yang, Z. L., Lu, G. M., Yang, G. F., & Zhang, L. J. (2017). PET/MR Imaging: New Frontier in Alzheimer’s Disease and Other Dementias. Frontiers in Molecular Neuroscience, 10, 343. 10.3389/fnmol.2017.00343

