## Supplementary Protocols for "Precise MRI-Histology Coregistration of Paraffin-Embedded Tissue with Blockface Imaging"

### Protocol for Precise Positioning of Camera, Light Source, and Microtome for Blockface Imaging

**Overview Setup:**

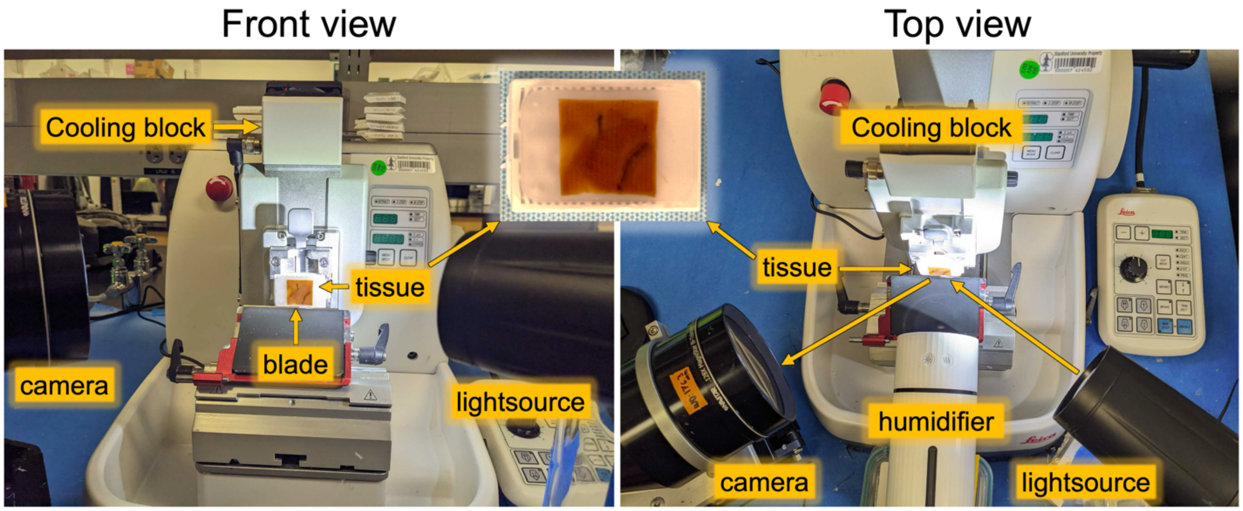


**Camera:** PL-D775 5.0MP rolling shutter CMOS USB3.0 color camera; Pixellink color sensor: MT9P006; mono sensor: MT9P031, Navitar, Inc.

**Bi-telecentric lens:** Bi-telecentric 0.0128X F/7 C-MOUNT lens, Magnification: 0.128X, Working Distance: 176.3 mm, Telecentricity: 0.05°, Field Depth: 31 mm, Average Transmittance from 460-630nm: 97%, Navitar, Inc.

**Microtome:** HistoCore NANOCUT R microtome, Leica, Inc.

**Cooling block**: Leica RM CoolClamp, Leica, Inc.

**Portable humidifier:** Palanchy, Inc.

### 1. Aligning the Microtome

1) Place a metal ruler flush against the backside of the microtome, ensuring it touches the tabletop to create a stable reference area for measurement.

2) Use an additional ruler to measure the distance between the edge of the microtome and the tabletop.

3) Confirm that the measured distance is consistent along the entire microtome edge to ensure proper alignment.

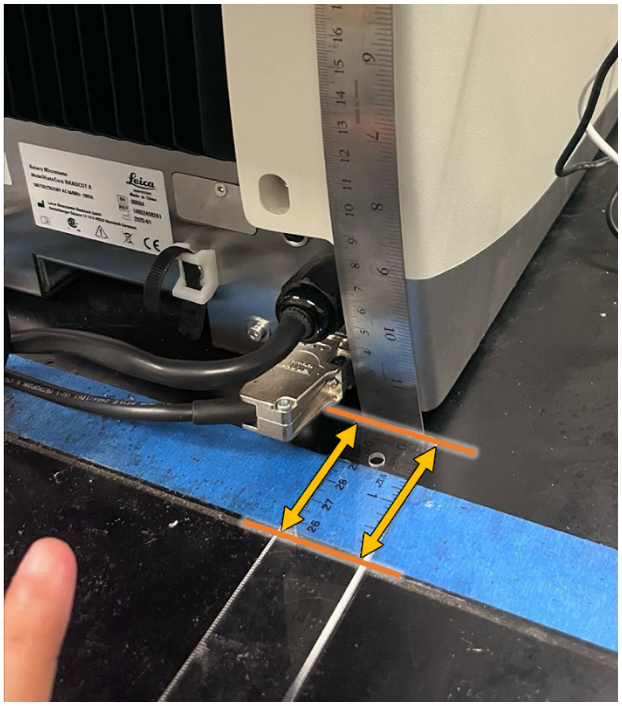

#### 2. Positioning the Knife Holder Base

1) Ensure that the edge of the knife holder base (upper arrow) is perfectly aligned with the edge of the microtome base (lower arrow).

2) Verify alignment visually and through manual measurements for precision.

3) Place the cutting angle at 5.

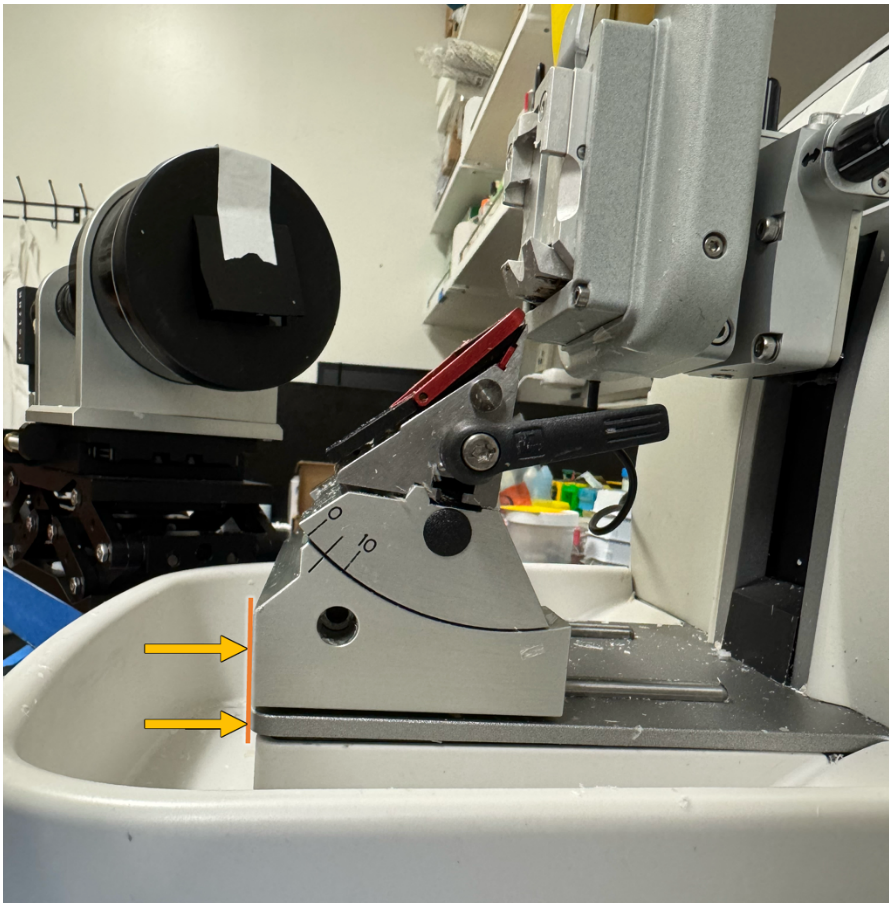

#### 3. Setting the Distance Between the Camera and Microtome

1) Position the block surface in the cutting position before measuring the working distance.

2) Place a ruler horizontally from the block holder lever to the tip of the camera lens cover.

3) Adjust the camera so the working distance between the lens and the middle line of the block surface should follow the working distance of the lens, 17.63 cm.

*4)* After accounting for the ~2-mm offset between the lens cap and the lens itself, we should measure approximately ~17.5 cm between the lens's front edge and the block surface's middle line.

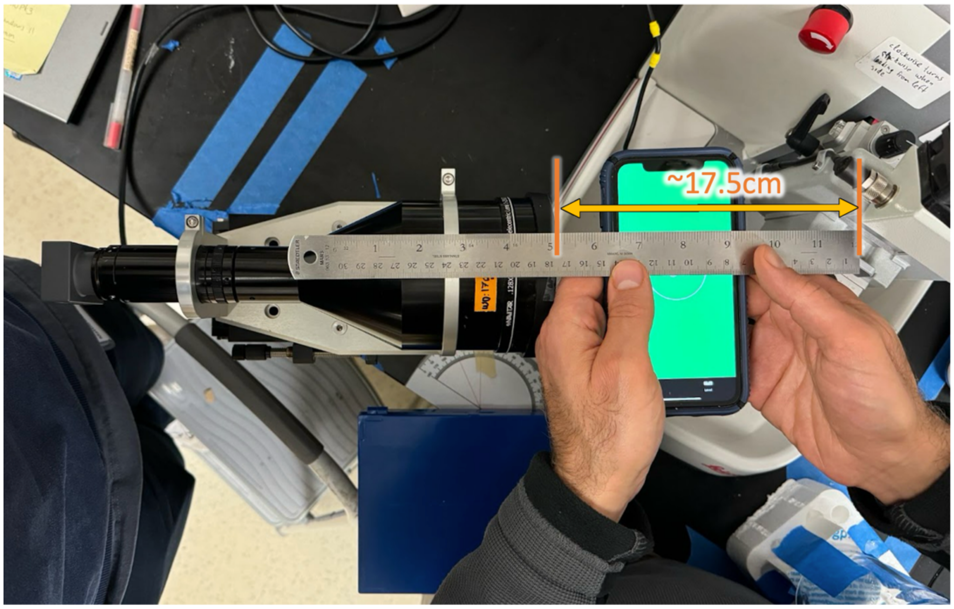

#### 4. Adjusting the Camera Angle

1) The camera should be positioned at Brewster’s angle (57° from the normal for the paraffin-air boundary).

2) Place a protractor near the camera to measure the angle between the camera and desk edges, ensuring it is set to 33°.

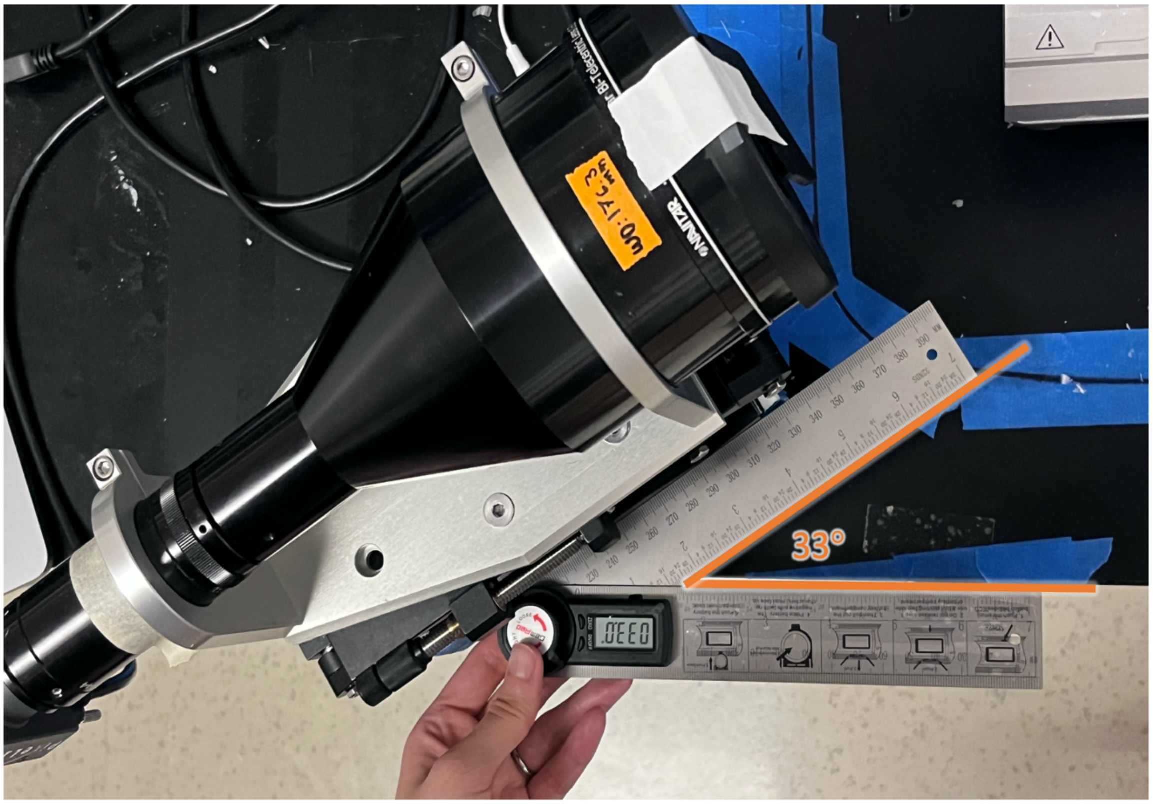

### 5. Light Source Positioning

1) Setting the Light Source at Brewster’s angle
    a. Adjust the light source to maximize brightness on the tissue block surface.
    b. Set the angle between the light source edge and the desk edge to 33°
    c. Identify Brewster’s angle by observing the inversion between gray and white matter and very bright reflective surface.
2) Transitioning from Brewster’s to the desired Brewster’s-Adjacent angle
    a. Increase the horizontal angle from 33° to 36°.
    b. Raise the light source by 2 cm.
    c. Tilt the light source downward by 1°.
    d. Make minor adjustments to ensure homogeneous illumination, the presence of some reflective lines, but not as many as at Brewster’s angle.
    e. Secure the light source.
3) Transitioning to Fully Off-Brewster’s Angle
    a. Further raise the light source by 5 cm.
    b. Tilt the light source downward by 17°.

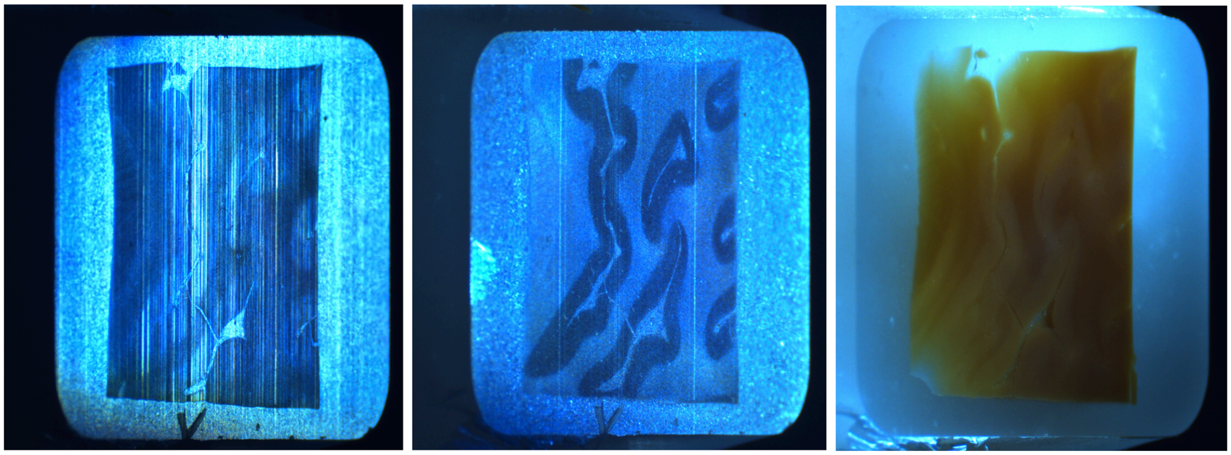

On Brewster’s    Brewster’s Adjacent    Off Brewster’s
